# Targeted modulation of protein liquid-liquid phase separation by evolution of amino-acid sequence

**DOI:** 10.1101/2020.10.20.347542

**Authors:** Simon M. Lichtinger, Adiran Garaizar, Rosana Collepardo-Guevara, Aleks Reinhardt

**Affiliations:** Department of Chemistry, University of Cambridge, Lensfield Road, Cambridge, CB2 1EW, United Kingdom; Department of Physics, University of Cambridge, Cavendish Laboratory, Maxwell Centre, JJ Thomson Avenue, Cambridge, CB3 0HE, United Kingdom; Department of Genetics, University of Cambridge, 20 Downing Place, Cambridge, CB2 3EJ, United Kingdom

## Abstract

Rationally and efficiently modifying the amino-acid sequence of proteins to control their ability to undergo liquid-liquid phase separation (LLPS) on demand is not only highly desirable, but can also help to elucidate which protein features are important for LLPS. Here, we propose an innovative computational method that couples a genetic algorithm to a sequence-dependent coarse-grained protein model to evolve the amino-acid sequences of phase-separating intrinsically disordered protein regions (IDRs), and purposely enhance or inhibit their capacity to phase-separate. We apply it to the phase-separating IDRs of three naturally occurring proteins, namely FUS, hnRNPA1 and LAF1, as prototypes of regions that exist in cells and undergo homotypic LLPS driven by different types of intermolecular interaction. We find that the evolution of amino-acid sequences towards enhanced LLPS is driven in these three cases, among other factors, by an increase in the average size of the amino acids. However, the direction of change in the molecular driving forces that enhance LLPS (such as hydrophobicity, aromaticity and charge) depends on the initial amino-acid sequence: the critical temperature can be enhanced by increasing the frequency of hydrophobic and aromatic residues, by changing the charge patterning, or by a combination of both. Finally, we show that the evolution of amino-acid sequences to modulate LLPS is strongly coupled to the composition of the medium (e.g. the presence or absence of RNA), which may have significant implications for our understanding of phase separation within the many-component mixtures of biological systems.

---

Liquid-liquid phase separation (LLPS) of multivalent biomolecules (e.g. proteins and nucleic acids) is an important mechanism employed by cells to control the spatio-temporal organisation of their many components.^1,2^ Biomolecular condensates, or membraneless organelles, such as stress granules,^3^ P-granules,^4,5^ the nephrin–NCK–WASP system^6^ and the nucleoli,^7^ are formed by LLPS and have diverse biological functions. LLPS inside cells plays a very diverse range of roles beyond membraneless compartmentalisation, such as in gene silencing via heterochromatin formation,^8–10^ in gene activation by facilitating the formation of super-enhancers,^11^ in buffering cellular noise, in modulating enzymatic reactions and in sensing pH changes in the skin.^14^ Moreover, aberrant LLPS is associated with the onset of various human pathologies, such as degenerative diseases and cancer.^15^ Biomolecular condensates have also recently been proposed as promising new tools to partition anti-cancer drugs preferentially to cancer cells.^16^ However, some biomolecular condensates emerge spontaneously inside cells without as-yet clearly identified functions; it has been hypothesised that some of these might be implicated in the emergence of phase-separation-related pathologies.^15^

The thermodynamics of phase separation is driven by the competition between interaction enthalpies and the entropic favourability of mixing.^17–19^ In the context of protein solutions, LLPS is principally driven by dipole–dipole, charge–charge, cation–π and π-stacking interactions,^20^ whose strengths are modulated by the experimental conditions, including the temperature, pH and salt concentration.^21^ In biomolecular systems, the multivalency in mixtures is thus the main physical parameter that defines the ability of a system to undergo LLPS:^6,15,22–25^ biomolecules with higher valencies can establish a larger number of weak attractive interactions with other species and hence form a more stable condensate.

The connections between LLPS and cellular function, and between aberrant LLPS and human pathologies, suggest that learning how to control or even revert the phase separation of proteins by subtly mutating their amino-acid sequences would be highly desirable. However, the sequence space of even the smallest naturally occurring proteins is immense, which makes the task of choosing mutations manually extremely inefficient; what is more, even if small-scale modifications of a single protein that promote phase separation might be possible to design manually with some physical intuition, biomolecular LLPS is a collective phenomenon involving many weak interactions, and it is not at all straightforward to anticipate how small sequence modifications affect the phase behaviour of a protein mixture without the use of an algorithm.

Indeed, optimising biological LLPS is especially difficult because *in vivo* biomolecular condensates are highly multicomponent systems.^23,26–28^ Furthermore, over 270 distinct multivalent proteins have been shown to undergo LLPS *in vitro*.^29^ Despite this complexity, the properties of condensates can often be approximated well by considering just a fraction of biomolecules, known as ‘scaffolds’, since such molecules tend to dominate the phase behaviour:^6,22,30^ the addition of biomolecules that are recruited to condensates via their interactions with scaffolds, termed ‘clients’,^6,22^ impacts the stability of condensates only marginally,^25^ making the problem somewhat more manageable.

A wide body of work has significantly advanced our understanding of how changes in amino-acid sequence transform the phase behaviour of different proteins.^24,31–36^ Notably, tightly integrated experiments and simulations with the ‘stickers and spacers’ model explain how changing the number and patterning of aromatic residues can alter the phase diagrams of prion-like-domain proteins in a predictable manner.^24^ Experiments also demonstrate that multimerising the arginine/glycine-rich RGG domain of LAF1 leads to controllable phase separation,^37^ and minimal coarse-grained simulations of point mutations of two designer proteins exemplify how on-demand modulation of protein phase behaviour can be achieved *in vivo*.^38^

Alongside globular domains, intrinsically disordered regions (IDRs) are thought to be one of the main drivers of LLPS in protein systems.^39,40^ Various theoretical approaches, including random-phase approximation theory, have been applied to LLPS of IDRs.^41,42^ Such treatments can rationalise aspects of charge distribution in phase-separating IDRs, and their relative simplicity has allowed us to gain significant intuition for the electrostatic aspects of phase separation. However, the scope of theory is limited by the inherent approximations that are of necessity made and the sheer complexity of biological multi-component systems. More realistic representations of proteins are usually too complex for analytical treatment, but can be studied in computer simulation, giving us molecular-level insight into the phase behaviour of biological systems. Computational approaches hold significant promise of enabling us to probe amino-acid sequence space efficiently and to design protein mutations which enhance or inhibit LLPS of a protein solution. Although all-atom simulations of LLPS in explicit solvents are only now slowly becoming computationally tractable^43,44^ due to the exponential scaling of search space with system size, theory and simulations using coarse-grained potentials – from patchy particles to lattice models^24,25,33,37,43,45–52^ – have shown that significant microscopic insight can be gained into the physics of phase separation.

Here, we develop a genetic-algorithm approach coupled to the sequence-dependent coarse-grained model of proteins with amino-acid resolution of Dignon *et al*^33^ [Fig. 1(a)(i)]. Our algorithm is anchored in a fitness function that is fast enough to be evolved and that represents a good proxy for the critical solution temperature, which measures the ability of a protein to phase-separate. With this approach, we systematically evolve the amino-acid sequences of the IDRs of three naturally occurring proteins that are known to phase-separate *in vitro* via homotypic interactions, and we show that we can drive the genetic algorithm either to enhance or to inhibit their LLPS. By shuffling the aminoacid sequences in chunks of varying lengths, we also identify the binding domains of the IDRs that are essential to drive LLPS (the ‘stickers’) and the connecting regions (the ‘spacers’).^24,34,49^ By probing LLPS in the vicinity of known phase-separating sequences, we can infer which features of a sequence drive phase separation in biological systems. While previously, artificial sequences have been probed in a systematic way, for instance in the context of charge patterning,^43^ our work also complements very recent results obtained on LAF1-IDR37 and Ddx4-IDR.^52^

**FIG. 1.**
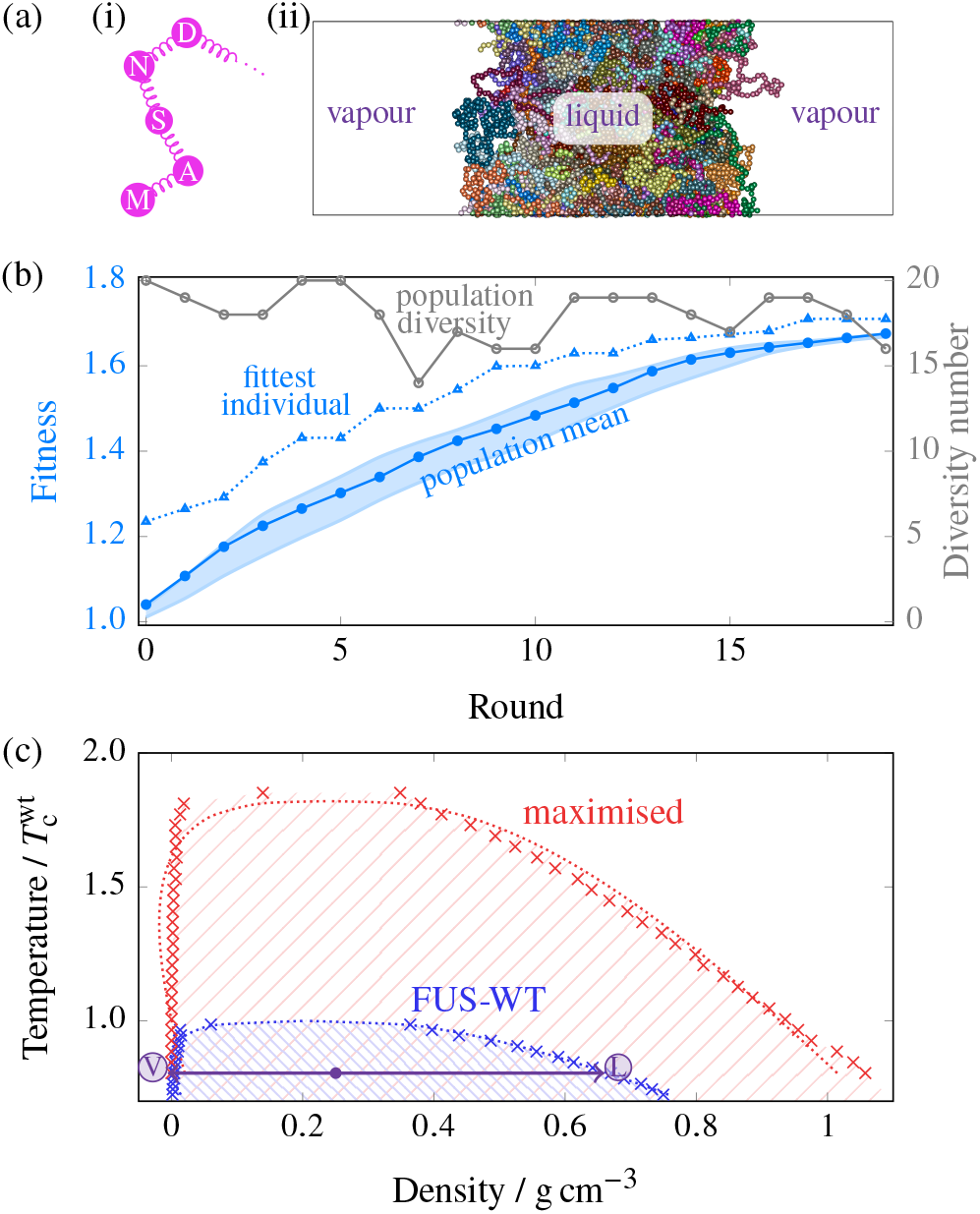
(a) (i) A schematic representation of the model used. Each amino acid is represented by a bead, and beads are connected with harmonic springs. (ii) A snapshot of a typical simulation cell exhibiting coexistence between a liquid-like (protein-rich, high-density) fluid and a vapour-like (protein-poor, low-density) fluid. The box is periodic in all directions. Different colours are used to represent beads in different protein chains. (b) Typical genetic-algorithm progression for FUS when maximising the width of the phase diagram. The fitness function [Eq. (2)] increases by ~65 % over 20 rounds. The fittest individual is 5 % to 20 % fitter than the mean in most rounds. The population diversity, i.e. the number of distinct sequences present in the overall population of 20, remains high throughout the run. The shaded area corresponds to the range of values of the mean fitness obtained from 3 independent genetic-algorithm runs. (c) Comparison of representative phase diagrams before and after genetic-algorithm runs, confirming that the fitness function choice was suitable. Pale hatched lines indicate the approximate region of phase separation for each case. Error bars in the density evaluations are smaller than the symbols, and dotted lines are fits as detailed in Sec. SI-S1-b. The point labelled in violet corresponds to the snapshot shown in panel (a), with the densities of the vapour-like and liquid-like fluids labelled ‘V’ and ‘L’, respectively. All densities reported are for the protein in a massless implicit solvent.

## I. PROTEIN PHASE BEHAVIOUR CAN BE GUIDED BY A GENETIC ALGORITHM

A free choice from amongst the 20 canonical amino acids in a protein with *n* residues amounts to an *n*-dimensional vector with 20^*n*^ possible sequences, where for each sequence one might attempt to compute some property that characterises the sequence’s LLPS behaviour. One possible quantity that can serve this purpose is the upper critical solution temperature *T_c_*, above which no de-mixing occurs. However, an exhaustive search of sequence space would be prohibitively expensive. Moreover, finding the ‘optimal’ critical temperature, however we might choose to define it, is not itself the aim. For example, a poly-F chain has a particularly high *T*_c_ (see Sec. SI-S2) in the protein model we have used, but studying it in detail is not particularly helpful in understanding what drives biological phase separation. We instead focus on biologically occurring proteins and evolve their amino-acid sequences with the aim of either extending or narrowing the range of thermodynamic conditions where homotypic LLPS occurs. In particular, we are interested in the effect of relatively small changes to the amino-acid sequence on phase separation, as these are instrumental to understanding how modifications can be designed to control the phase behaviour of proteins *in vivo*. Moreover, such modified sequences might more easily be introduced into cells.

To determine sets of mutations that shift the phase behaviour of a protein, we start from a reference amino-acid sequence and perform direct-coexistence molecular dynamics (MD) simulations of a sufficiently large number of copies of that protein (see Fig. 1(a)(ii) and Sec. SI-S1-b). In direct-coexistence simulations, we explicitly simulate two different phases – a protein-enriched solution in contact with a protein-depleted solution – in contact with each other in the same simulation box. By performing such simulations at several temperatures, we can approximate the compositional phase diagram, which indicates which phase is thermodynamically stable as a function of temperature and protein density. We then seek to evolve the protein towards enhanced or inhibited LLPS by developing a genetic algorithm (see Methods) that iteratively proposes stochastic amino-acid sequence mutations, selecting a few at each iteration amongst those that induce the strongest effect in the protein LLPS, and mutating again. Whether such a genetic-algorithm approach can succeed in evolving protein phase behaviour depends on the quality and efficiency of the fitness function used to control it. Our genetic algorithm uses the difference in composition densities of the protein-poor and protein-rich phases at constant volume as its fitness function. As we show below, such a function is both computationally inexpensive and a good metric to determine whether a set of mutations would result in enhanced or inhibited LLPS. Although the critical solution temperature of a protein mixture may seem like an obvious order parameter to determine whether a specific set of mutations promotes or inhibits LLPS – by raising and lowering the critical solution temperature, respectively – computing it in every round of the evolution process when using a residue-resolution coarse-grained protein model is computationally infeasible. This is because estimating the critical solution temperature requires an evaluation of a full phase diagram of the protein solution, which in turn requires either the use of very expensive free-energy methods,^53^ or performing direct-coexistence simulations at a number of different temperatures, each involving long MD simulations of a large number of copies of the same protein, and analysing the results to extrapolate the data and estimate the critical temperature. By contrast, evaluating the difference in composition densities requires only one set of direct-coexistence simulations to be run at a fixed sub-critical temperature (i.e. below *T*_c_).

## II. THE CASE OF THE PLD OF FUS

### A. The range of stability of LLPS can be evolved

As an initial model system, we investigate the behaviour of the prion-like domain (PLD) of the FUS protein, an IDR mostly devoid of charged residues. Although the PLD of FUS only phase-separates *in vitro* at somewhat extreme conditions with respect to the physiological ones (namely low salt concentrations of 37.5 mM NaCl and high protein concentrations of 6 μM to 33 μM),^54^ PLD-PLD interactions and PLD-arginine-rich domain interactions drive LLPS of the full FUS protein under physiological conditions both *in vitro* and in cells.^55^ We use direct-coexistence molecular dynamics simulations to approximate the compositional phase diagram (i.e. in the temperature versus protein-density space) of the PLD of FUS. We then seek to evolve the system’s critical solution temperature by introducing our genetic algorithm (see Methods), which allows us to mimic, broadly speaking, the evolutionary pathways that might drive phase separation in nature. Starting from the reference aminoacid sequence of FUS PLD (WT-FUS) given in Sec. SI-S9, we use our genetic algorithm to attempt separately to maximise and to minimise the width of the binodal curve of the compositional phase diagram. We show the population fitness [Eq. (2)] and the diversity of the population as functions of the genetic-algorithm round in Fig. 1(b) for the maximising case; analogous results for the minimising case are shown in Sec. SI-S3. The genetic algorithm is effective in increasing the population fitness in each case, and in both cases the population diversity remains high, indicating that no premature convergence occurs. In Fig. 1(c), we show that the phase diagram of the evolved FUS PLD for the maximising case exhibits a large increase in the range of temperatures and densities at which LLPS occurs, with the critical temperature increasing by ~65 % compared to the WT FUS PLD. Although we have used the width of the phase diagram as a proxy for the critical solution temperature, Fig. 1(c) confirms not only that the critical solution temperature behaves in the expected way, but also that genetic algorithms with simple fitness functions can significantly perturb the LLPS behaviour, leading to an effective gradient in sequence space. These results suggest that a genetic algorithm can be used to search the sequence space of proteins efficiently and can help identify those sequence mutations that yield meaningful changes in the proteins’ compositional phase diagrams.

### B. LLPS evolution can be driven by changes in hydrophobic and aromatic residue composition

We analyse the extent to which all interactions other than direct charge–charge interactions, which here we term collectively as ‘hydrophobicity’, govern the evolution of the phase behaviour of proteins by estimating each amino acid’s relative degree of hydrophobicity. We use the hydrophobicity scale proposed in Ref. ^56^, which is quantified as the *λ* parameter in the coarsegrained model of Dignon *et al.*,^33^ and which can be used to scale the well depth of the modified Lennard-Jones potential in an amino-acid-specific way (see Sec. SI-S1-a). In Fig. 2(a), we show the amino-acid compositions of the maximised and minimised populations, broken down by amino acid and ordered by the extent of hydrophobicity, alongside the reference WT sequence.^57^ The amino-acid sequences of the WT, and examples of the minimised and the maximised FUS PLDs are given in Sec. SI-S10. In the maximised case, there is a general shift towards higher hydrophobicity, whilst the minimised case shows a trend towards highly polar and charged amino acids. These trends in amino-acid composition confirm that, even though there are more strongly hydrophobic than weakly hydrophobic amino acids available for insertion, evolution of the FUS PLD is able to be driven in both the hydrophobic and hydrophilic directions, illustrating the robustness of the genetic-algorithm approach.

**FIG. 2.**
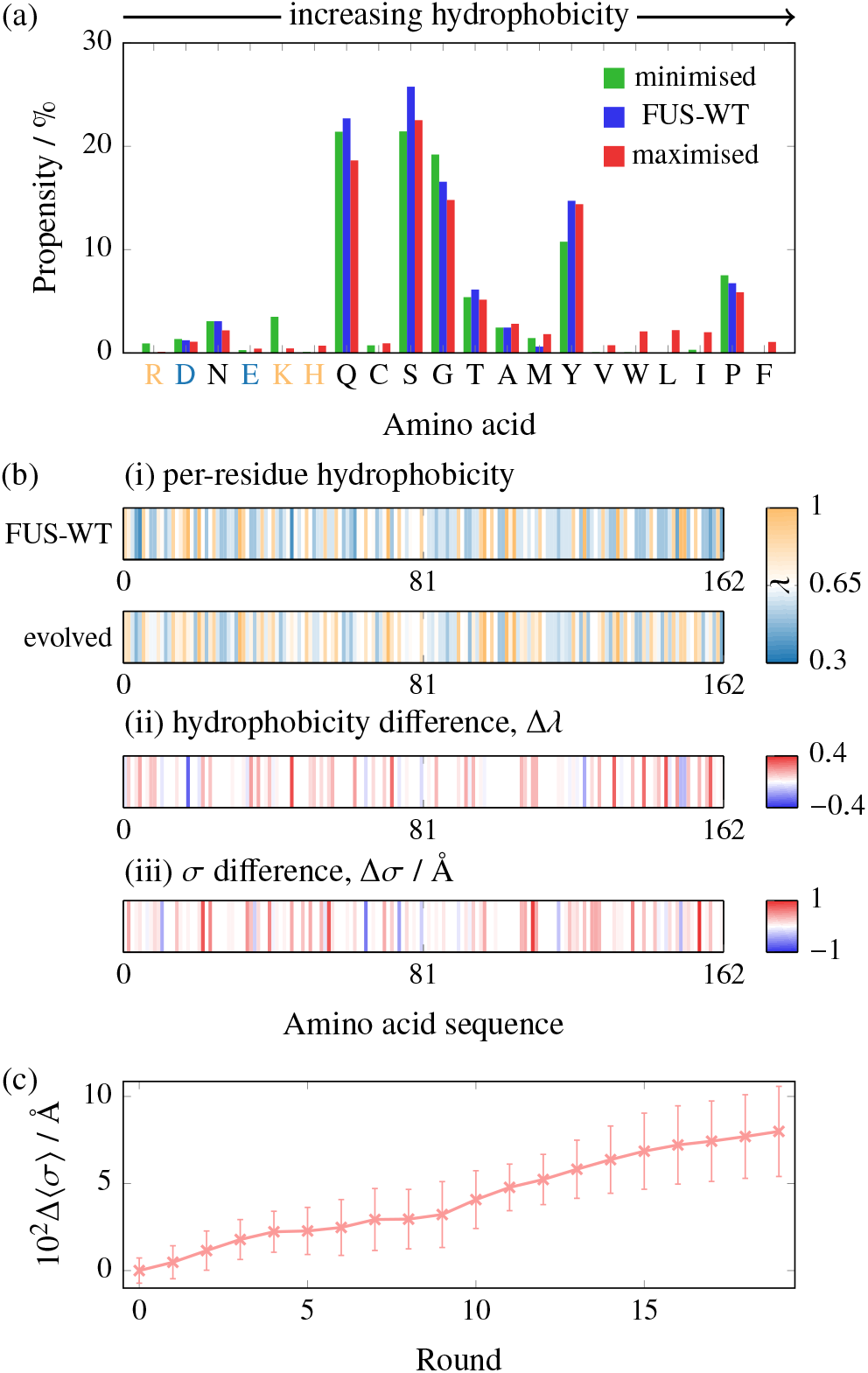
(a) Amino-acid composition before and after maximising and minimising the phase diagram width starting from WT-FUS. Amino acids are plotted in order of increasing hydrophobicity [see Table S1]. Positively charged amino acids are indicated in light orange and negatively ones in light blue. In maximising genetic-algorithm runs, hydrophobic amino acids become favoured, whilst the converse holds for minimising genetic-algorithm runs. (b) Map of (i) the per-residue hydrophobicity along the sequence, (ii) the change from the wild type to the evolved protein after maximisation and (iii) the change of the per-residue amino-acid size (*σ*). The data for the evolved protein are averaged over the entire population at the end of 3 independent genetic-algorithm runs. No larger-scale regional preference for modification is readily apparent. Trends in hydrophobicity and *σ* are largely correlated. (c) The average *σ* value of the amino acids in the population increases over the course of the genetic-algorithm run. Error bars are standard deviations of the averaged *σ* value of individual sequences with respect to the pooled population from the three genetic-algorithm runs.

In addition to the attractiveness of the hydrophobic interactions, a further factor determining the strength of hydrophobic interactions is the size of each amino acid. We quantify this by *σ*, the Van der Waals radius of each amino acid (see Table S1). In Fig. 2(c), we show that the average size of the amino acids in the sequence population increases as a function of the genetic-algorithm round, implying that the average size of the amino acids increases through evolution. Furthermore, as shown in Fig. 2(b) (iii), although the effects on hydrophobic attractiveness and *σ* values largely correlate at most residues of the protein sequence, this is not invariably the case, giving us the first indication that the size of the amino acids could play an independent role in determining LLPS properties. Although the overall increase in amino-acid size might be explained at least in part by the amino-acid size defining the range of the hydrophobic attractions, we hypothesise that the main physical driving force explaining the increase of both hydrophobicity and size is the ability of larger and more hydrophobic amino acids to form a more densely connected, and in turn more stable, condensed liquid-like protein-rich phase.^25^ To test this hypothesis, we compute the radial distribution function of the protein-rich phase for both FUS-WT and a maximised sequence at a common number density (Fig. S4). This pair correlation function exhibits a more pronounced nearest-neighbour maximum, indicating a greater degree of local structure and an increase in the number of nearest-neighbour beads compared to the WT, which has previously been shown^25^ to correspond to a greater protein valency.

Because the sequence of the WT PLD of FUS only contains two (negatively) charged amino acids, its LLPS must be driven by hydrophobicity. However, our results show that the FUS sequence lies on a hydrophobic gradient in sequence space: an increase in hydrophobicity effects an increase in the critical solution temperature. It has been proposed^58,59^ that the driving force for the LLPS of this FUS IDR is specifically the interactions between tyrosine residues dispersed through the sequence. Although such interactions are only implicitly captured in the coarse-grained protein model we use through its hydrophobicity parameter, and thus the distribution of amino acids obtained follows broad trends rather than converging to a distinct amino acid or motif, our results are consistent with previous work,^58,60,61^ and meaningful trends in composition and sequence can be observed from our simulations. Our results thus suggest that evolving a protein sequence which is dominated by hydrophobic residues, as is the case for the PLD of FUS, towards enhancing its propensity for LLPS is efficiently achieved by protein mutations that increase the average attractiveness and size of the protein’s uncharged residues.

### C. Hydrophobic patterning has a minimum length scale

In Fig. 2(b), we show how the accumulated changes in sequence, represented as the hydrophobicity of a residue, map onto the WT-FUS sequence. There are no larger-scale regions along the sequence where modifications occur preferentially; instead, there appears to be a stochastic increase in hydrophobicity, with less hydrophobic residues being replaced by more hydrophobic ones. However, since short runs of the genetic algorithm cannot result in perfectly uniform replacement attempt probabilities, we cannot expect to be able to resolve small-scale features in amino-acid sequence space. We have therefore first replaced successive chunks of the amino-acid sequence with a fixed amino acid. The conventional approach to probe the function of specific residues is alanine scanning.^62–64^ As we are interested in how hydrophobicity affects phase behaviour, in this case, we mutate amino acids to glycine rather than alanine, as the former has the median hydrophobicity in the coarse-grained protein model (see Table S1). Although in experiments or all-atom simulations, such a replacement may be less appropriate, as glycine disrupts protein secondary structure by its dihedral angle preferences,^65^ in the CG model this effect is immaterial, as no conformational terms are considered. Fig. 3(a) shows the results of a glycine scan in sequential chunks of 6 amino acids, projected onto the chunk-averaged hydrophobicities of the WT protein. The curves anti-correlate for most of the sequence, reflecting that in hydrophobic stretches, mutating to glycine decreases hydrophobicity and thus decreases fitness, while the converse holds for hydrophilic stretches.

**FIG. 3.**
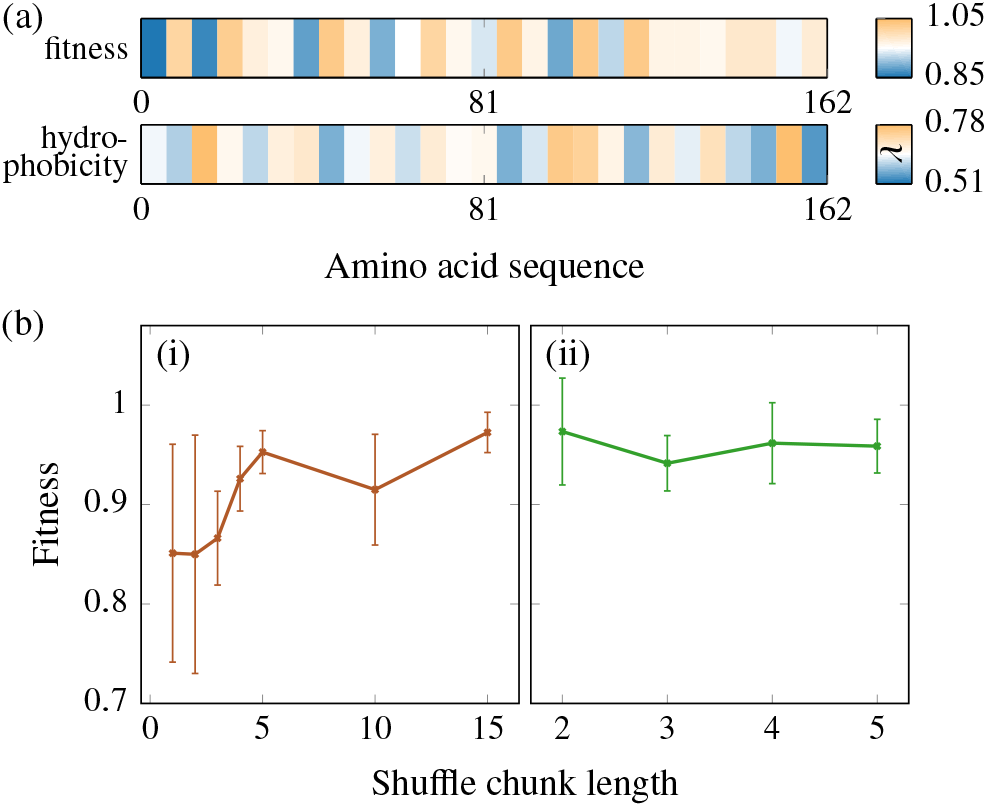
(a) The resulting value of the fitness function (relative to the WT fitness), when chunks of 6 amino acids of FUS-WT are separately replaced with glycine, and the average hydrophobicity of these same 6-residue chunks of FUS-WT, with colours distributed about the hydrophobicity parameter of glycine (λ(G) = 0.649). Where glycine represents a gain in hydrophobicity, the fitness change is largely positive, and vice versa. (b) (i) The fitness of the system as a function of the chunk length following the shuffling of approximately 100 amino acids. Error bars show the standard deviation across several shuffling runs. There is a significant difference in the fitness and the error bars for small chunks of 1 and 2 amino acids, whereas the error bars for the larger chunk sizes are considerably smaller. (ii) Analogous results for shuffling runs with a hydrophobicity bias, where exchanges were allowed only amongst the top 30 % of chunks by hydrophobicity. Representative fitness functions as a function of the number of amino acid pairs shuffled are shown in Fig. S5.

In order to probe the resulting sequence further, we ‘shuffle’ the sequence without changing its overall amino-acid make-up. We choose chunks of varying lengths by randomly choosing two positions along the sequence and exchanging chunks of *l* amino acids starting from those two positions. The ends of the sequence are treated periodically to ensure no positional selection bias against the ends. We record the fitness function of the protein sequences as a function of the total number of amino-acid pairs changed, i.e. the number of exchange steps multiplied by *l*. In Fig. 3(b), we show the variation of the fitness with chunk length after ~100 amino acids have been shuffled. The error bar, which shows the standard deviation across several independent runs, is a useful measure of the sensitivity of the fitness function to amino-acid sequence. Very small chunk lengths, particularly of 1 or 2 amino acids, are highly disruptive to phase separation, while larger chunk lengths only cause smaller modulations. From these results, we can conclude that segments of 2-3 successive amino acids are crucial in driving LLPS in the PLD of FUS, representing the length scale of some sequence feature. To investigate the nature of this feature, we repeated the shuffling analysis with a hydrophobicity bias, where only the most hydrophobic of all possible contiguous chunks are exchanged. The dependence on chunk size largely disappears, implying that it was small hydrophobic patches that were previously disrupted by shuffling (Fig. 3(b)(ii)). The phase separation therefore appears to be governed by a hydrophobic patterning of a minimum length scale of 2-3 amino acids. This is consistent with the ‘stickersand-spacers’ paradigm of phase-separating proteins, in which proteins are considered to comprise stickers – corresponding to the attractive protein regions that drive LLPS, in our case the small chunks of 2-3 amino-acid residues – that are connected by less attractive regions termed spacers.^24,34,49^

## III. CHARGE PATTERNING MAY BE AN ALTERNATIVE DRIVING FORCE FOR EVOLUTION OF PROTEIN PHASE BEHAVIOUR

Not all proteins that exhibit LLPS are expected to be governed by the same driving force. For example, the patterning of charges has been suggested to contribute to LLPS in charge-rich proteins,^42,43^ while the phase separation of the intrinsically disordered region of the protein hnRNPA1, which belongs to the family of heterogeneous nuclear ribonucleoproteins, has been suggested to be driven by the interaction between linearly dispersed aromatic residues within the polar sequence.^24,32^

Here, we will use the hnRNPA1-IDR as a test case to complement the behaviour observed for FUS. We define the IDR of hnRNPA1 as its first 135 residues, 11.9 % of which carry a formal charge, and which has been shown to phase-separate *in vitro*.^54^ We used a genetic-algorithm-driven evolution of the same fitness function as in the case of FUS; the genetic-algorithm approach results in an increase of fitness whilst maintaining the population diversity [Fig. S6(a)], and the increased fitness function is again a successful proxy for the upper critical solution temperature [Fig. S6(b)]. However, the change in amino-acid composition upon applying the genetic algorithm is qualitatively different from the hydrophobicity-driven case of Subsect. IIB, as hydrophilic/charged residues are not lost and hydrophobic residues appear, statistically replacing some of the highly abundant amino acids of intermediate hydrophobicity [Fig. 4(a)], indicating that the driving force for phase separation may be different from the case of FUS-PLD.

**FIG. 4.**
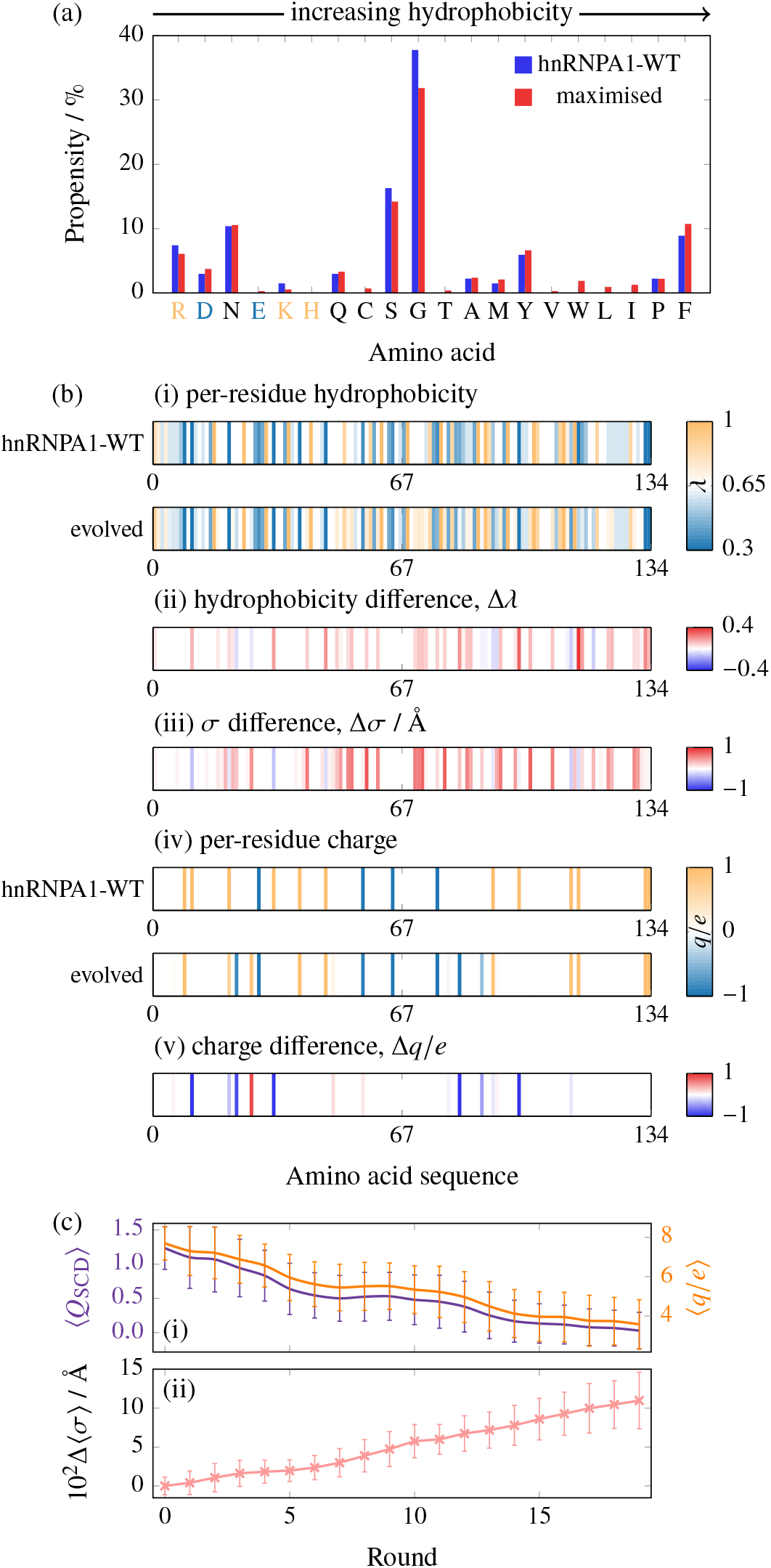
(a) Amino-acid composition before and after maximising the phase diagram width for hnRNPA1, revealing the appearance of hydrophobic amino acids while no bias against charged or polar amino acids is observed. Amino acids are plotted in order of increasing hydrophobicity [see Table S1]. Positively charged amino acids are indicated in light orange and negatively ones in light blue. (b) Map of (i) the per-residue hydrophobicity along the sequence, (ii) the change from WT hnRNPA1 to the evolved protein, (iii) the per-residue change *σ* value and (iv), (v) analogous maps for the charge. The data for the evolved protein are averaged over the entire population at the end of 3 independent genetic-algorithm runs. Partial charges reflect only partial carriage in the population. Some charges have appeared and some have disappeared; the overall balance is towards charge neutralisation. (c) (i) The sequence charge decoration (SCD) and the charge number (*q/e*) of the population over the course of the genetic-algorithm run, indicating an evolutionary charge neutralisation. (ii) The average *σ* value of the amino acids in the population increases over the course of the genetic-algorithm run. Error bars are standard deviations of the averaged *σ* value of individual sequences with respect to the population.

To investigate this further, we have analysed the initial and final populations in the genetic-algorithm runs. As in the case of the PLD of FUS, the hydrophobic attractiveness and the average size of amino acids of the IDR of hnRNPA1 increase to raise the propensity for LLPS (Fig. 4(b)(i-iii)), and as in the case of FUS, the per-residue effects of hydrophobic attractiveness and amino-acid size are largely, but not fully, correlated.

### A. Charge patterning and hydrophobicity can co-evolve…

Additionally, there is a substantial difference in terms of charge patterning over the course of the genetic-algorithm run [Fig. 4(b)(iv),(v)]. Charges are both created and lost across the sequence, but not uniformly so. We show in Fig. 4(c)(i) two measures of the charge patterning, the net charge of a protein chain, and the sequence charge decoration (SCD), defined as^66^

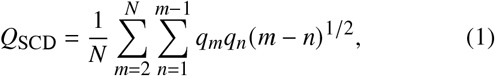

where *q_i_* is the formal charge number of residue *i* and *N* is the length of the amino-acid sequence. *Q*_SCD_ has been shown to anti-correlate with the upper critical solution temperature of an IDR.^43^ These two parameters show a virtually identical evolution through the genetic-algorithm run, which indicates that, in this case, the decrease in charge separation as measured by *Q*_SCD_ results from a net decrease in the overall charge of the protein. Specifically, this arises from the creation of a larger number of negative than positive charges.

A considerable amount of work has already been done in the context of the role of charge patterning.^37,42,67–70^ Although it is perhaps not overly surprising that a more even distribution of positive and negative charges allows for the largest number of attractive interactions, which in turn drives the formation of liquid-like phases, we show in Section IV that the precise nature of the optimal sequence depends on the medium in which the proteins of interest exist.

The local gradient in sequence space around hnRNPA1-WT has components in both hydrophobicity and charge redistribution. However, in the literature, hnRNPA1 LLPS is not commonly associated with charges.^24,32^ This prompts the question of how important the factors are in absolute terms. A crude estimate can be obtained from an analysis of the components of the pairwise energy in our simulations, shown in Table I. In particular, we split up the energy into a ‘hydrophobic’ (LJ) contribution and a Coulomb (electrostatic) contribution. Both components become more favourable over the course of a genetic-algorithm run, indicating that the sequence-space gradient towards higher *T*_c_ encompasses both charge rearrangement and hydrophobicity. These effects operate in parallel, but the hydrophobicity component contributes considerably more to the attractive energy in absolute terms.

### B. …but need not necessarily, even in charge-rich sequences

To check the applicability of the two mechanisms driving the evolution of the capacity to undergo LLPS that we have identified to other protein sequences, we have also investigated an IDR of the protein LAF1, which is a DDX3-family RNA-helicase found enriched in *C. elegans* P-granules, in which it drives phase separation. The IDR we have focussed on has been shown to be both necessary and sufficient for LLPS.^71^ It contains a significant proportion of charged amino acids, with 22.4 % of its 168 residues carrying a formal charge. This IDR has also been shown to phase-separate in simulations of the CG model used here,^33^ and a recent study^37^ combining CG simulations, all-atom simulations and turbidity assays has identified a sticky hydrophobic stretch as well as tyrosine and arginine residues to be involved in LAF1 LLPS. Additionally, it has been suggested^37^ that the even distribution of charges across the sequence may suggest that charge patterning is a controlling determinant of LLPS. Simulations and *in vivo* experiments have been carried out in corroboration of this hypothesis.^37^

In order to compare the behaviour of LAF1 to the two cases already considered, we have evolved its sequence using the same genetic algorithm. As before, we have computed the phase diagram at the end of the genetic-algorithm run. Although the genetic-algorithm progression is slower with this system and finite-size effects are more pronounced, the genetic algorithm with this fitness function can successfully increase the critical solution temperature [Fig. S7]. Compared to both FUS and hnRNPA1, the composition of the resulting optimised sequence population is less significantly changed [Fig. 5(a)], although this is consistent with the fact that the fitness function increases more slowly and the overall critical solution temperature is only ~30 % higher than the wild type in the simulations considered (compared to 65 % and 50 % for FUS and hnRNPA1, respectively). Nevertheless, there is a limited increase in hydrophobicity [Fig. 5(a-b)], with no region particularly favoured in terms of increased hydrophobicity, even though the amino acids early in the sequence are on average more hydrophobic than later ones. The change in the range of hydrophobic interactions, however, as quantified by the average *σ* values (Fig. 5(c)), is more significant (*Δσ* = 0.0396 Å over 10 rounds). This is comparable to the level of change we observed in FUS (*Δσ* = 0.0322 Å after the first 10 rounds). In particular, almost all changes made to the sequence by the genetic algorithm lead to larger amino-acid sizes, even though in terms of hydrophobic attractiveness their effects are much more varied. This leads us to speculate that the extended range of attractive interactions may be the dominant factor in driving the evolution of hydrophobic interactions in this case, rather than the *λ* values, which change less.

**FIG. 5.**
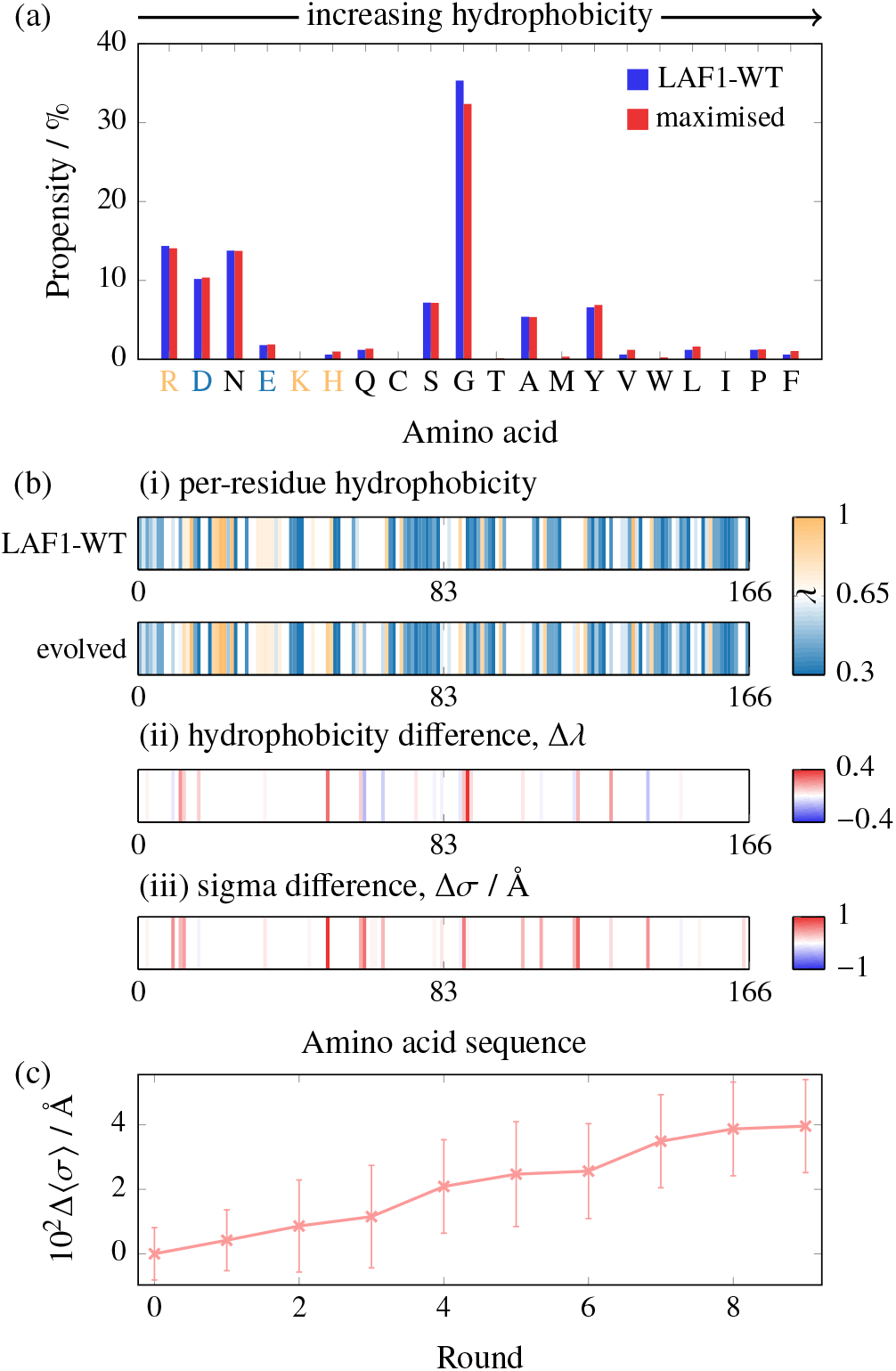
(a) Amino-acid composition before and after maximising the phase diagram width for LAF1. There is a slight general increase in hydrophobicity, whilst hydrophilic and charged residues are largely conserved. Amino acids are plotted in order of increasing hydrophobicity [see Table S1]. Positively charged amino acids are indicated in light orange and negatively ones in light blue. (b) Map of (i) the per-residue hydrophobicity along the sequence and (ii) the change from WT LAF1 to the evolved protein, and (iii), (iv) analogous maps for the charge. Data are shown for one genetic-algorithm run. Partial charges reflect only partial carriage in the population. There is a slight overall increase in hydrophobicity across the sequence, and there are more charges lost than created during the course of the genetic-algorithm runs. As opposed to the hydrophobicity, the *σ* values increase for almost all those residues that were changed by the genetic algorithm. (c) The average *σ* value of the amino acids in the population increases over the course of the genetic-algorithm run. Error bars are standard deviations of the averaged *σ* value of individual sequences with respect to the population.

The change in charge [Fig. S7(c)] is also relatively modest, and mainly entails the loss of existing net charge [Fig. S7(d)]. This is consistent with charge segregation, as quantified by the sequence charge decoration parameter, also shown in Fig. S7(d), which similarly decreases over the course of the genetic-algorithm run, but whose decrease is significantly less pronounced than in the case of hnRNPA1.

Similarly to the case of hnRNPA1, we show in Table I the change in the components of the average interaction energy for LAF1 before and after optimisation. However, in the case of LAF1, within the final evolved population, different sequences score rather differently in this analysis. Values for two evolved variants, termed LAF1*[a]* and LAF1*[b]*, are shown in the table. LAF1*[a]* behaves similarly to hnRNPA1, with both hydrophobic and Coulomb interactions interactions more favourable in the evolved sequence than in the wild-type, whilst in LAF1*[b]*, the Coulomb energy actually becomes less favourable. Since the genetic algorithm produces sequences in which the Coulomb attractions are enhanced as well as sequences in which they are weakened, charge patterning does not appear to act as an evolutionary driving force in LAF1. Based on these data we conclude that the evolution of the phase behaviour of the LAF1-IDR is primarily driven by hydrophobicity, and in particular by the size of the amino acids in the sequence.

**TABLE I.**
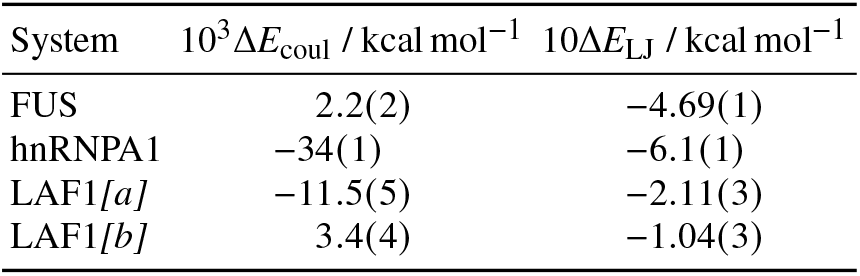
Changes in contributions to the average pairwise energy per bead between the WT and an evolved sequence of FUS, hnRNPA1 and LAF1. Standard deviations for the simulation averages are given in brackets and apply to the least significant figure. For LAF1, the evolved population is diverse in terms of these changes, and two representative examples are shown, labelled *[a]* (fitness 1.34) and *[b]* (fitness 1.28). For FUS, sequence evolution results in a change almost exclusively to the hydrophobic part of the pairwise energy. For hnRNPA1 and LAF1*[a]*, both Coulomb and hydrophobic interactions are more favourable in the evolved sequences, but hydrophobic interactions contribute more in absolute terms. For LAF1*[b]*, the Coulomb energy is less favourable in the evolved sequence. All data presented here are obtained at a simulation temperature of 200K, corresponding to 0.8 *T*_c_(FUS). While the overall average energies themselves depend significantly on temperature, the differences between WT and evolved sequence energies are largely independent of temperature in the range of interest.

| System | $10^3 \Delta E_{\text{coul}} / \text{kcal mol}^{-1}$ | $10 \Delta E_{\text{LJ}} / \text{kcal mol}^{-1}$ |
| --- | --- | --- |
| FUS | 2.2(2) | -4.69(1) |
| hnRNPA1 | -34(1) | -6.1(1) |
| LAF1[a] | -11.5(5) | -2.11(3) |
| LAF1[b] | 3.4(4) | -1.04(3) |

## IV. SEQUENCE EVOLUTION DEPENDS ON THE COMPOSITION OF THE MEDIUM

Inside cells, proteins are never isolated, and LLPS in multicomponent systems can be significantly different from that in single-component ones.^25^ It is therefore instructive to examine how genetic-algorithm driving behaviour changes upon adding a second component to the medium. To this end, we replace one eighth of the chains in the system with a poly-E protein of the same length as the respective protein of interest, but where all the amino-acid residues are glutamate (E). Such a poly-E protein has a high negative charge density, which gives it biophysical properties different from the protein it is replacing, making this a good initial test case for more complex multi-component systems.

We show in Fig. 6(a) the change in amino-acid composition at the end of the genetic-algorithm run in systems with and without the additional poly-E component, contrasted for (i) FUS and (ii) hnRNPA1. For FUS, the amino-acid composition change in the presence of a poly-E component is significantly different from the single-component case. In particular, in addition to the broad increase in hydrophobic residues that we have previously identified, in the presence of poly-E, positively charged residues become more prevalent, resulting in an overall positive mean charge of the sequences at the end of the run [Fig. 6(b)(i)]. The addition of poly-E affects the evolution of hnRNPA1 less significantly; however, we can still observe in Fig. 6(b)(ii) that the presence of poly-E slows down the loss of net charge over the course of the genetic-algorithm run. The higher initial positive charge content of hnRNPA1 already provides a greater electrostatic attractive energy with the additional poly-E component at the outset, reducing the driving force for the formation of positively charged residues compared to FUS. These results illustrate the robustness of our genetic-algorithm approach: we can not only probe the evolutionary driving forces resulting from changing the composition of the medium in which LLPS occurs, but can also gain insight into how such driving forces depend on the sequence of the protein of interest.

**FIG. 6.**
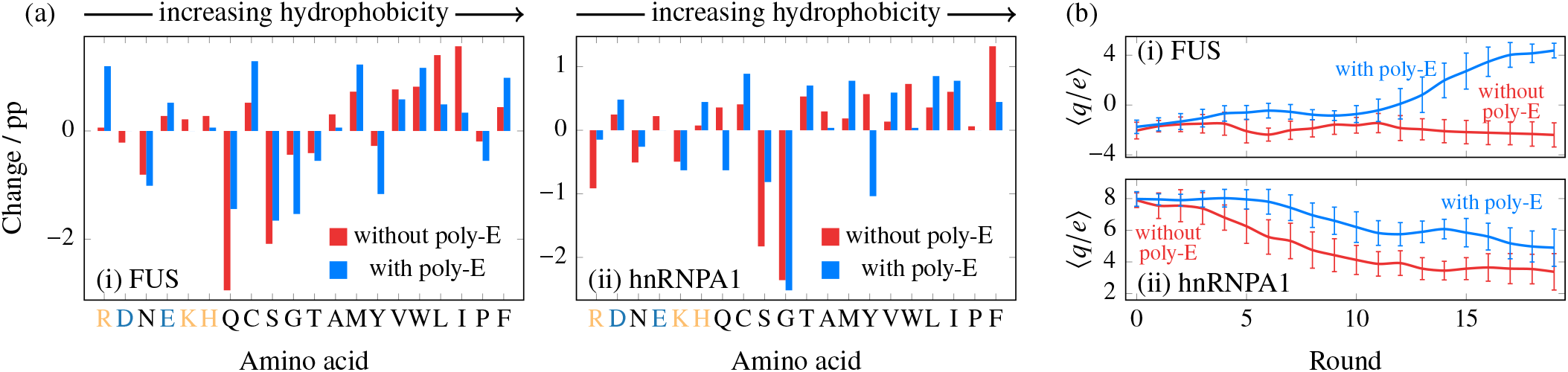
(a) Percentage-point difference in amino-acid composition after maximising the phase diagram width for (i) FUS and (ii) hnRNPA1. Amino acids are plotted in order of increasing hydrophobicity [see Table S1]. Positively charged amino acids are indicated in light orange and negatively charged ones in light blue. In each case, the ‘without poly-E’ data show results for the system optimised in the same way as before, and correspond to Fig. 2(a) and Fig. 4(a), respectively. The ‘with poly-E’ series corresponds to a system where 1/8 of the chains of the system are replaced with a poly-E protein that has been shown to mimic RNA. (b) Charge content as a function of genetic-algorithm round for (i) FUS and (ii) hnRNPA1. Error bars give standard deviations for the population at each round. The addition of poly-E changes the evolution behaviour significantly, particularly in the case of FUS, favouring higher charge content with the creation of new positive charges. In the case of hnRNPA1, changes in composition are less significant; however, the loss of positive charge content is slowed by the addition of poly-E. The evolutionary driving force thus depends not only on the initial composition of the protein, but also on the medium around a protein.

Interestingly, it has been shown that nucleic-acid chains can be represented as self-avoiding polymers;^72^ we can therefore interpret the poly-E sequence as a (relatively crude) mimic of RNA, a negatively charged polymer with no secondary structure. This may approximate intrinsically disordered single-stranded RNA with a short persistence length, such as poly(U),^73^ relatively well. Although RNA–protein interactions include cation–π interactions,^74^ which are not explicitly modelled in the coarse-grained potential used here, the charge-based driving force we observe suggests that the recruitment of phase-separating proteins into RNA-rich regions (and vice versa) could be driven by the pairing of opposite charges between the components of the mixture. Since all IDRs studied in this work are derived from proteins which in their full-length variants bind to RNA, and RNA itself can promote LLPS of proteins,^54^ this preliminary result is thus particularly exciting.

## V. CONCLUSIONS

In this work, we have proposed an efficient computational method to evolve naturally occurring phase-separating protein sequences. Evolving such sequences can provide insight into which sequence features drive LLPS, both when the proteins are in pure form and when they form part of a multi-component mixture; moreover, our approach could also be extended to design experimentally testable amino-acid sequence mutations which either inhibit or promote the LLPS of protein solutions. Our approach combines state-of-the-art molecular simulations of protein condensates, where each protein is described at the single-amino-acid resolution, with a genetic algorithm grounded in a new fitness function – the difference in composition densities of the protein-poor and protein-rich phases at constant volume – which is both a good proxy for the critical solution temperature and computationally far more tractable to obtain than the critical temperature itself. We have shown that such a simple and computationally inexpensive fitness function is sufficient to evolve the amino-acid sequences of naturally occurring proteins and to shift their phase behaviour in the direction we choose. Moreover, by analysing the effects of small changes to naturally occurring amino-acid sequences, we can draw conclusions about the molecular origins of the local gradient in amino-acid sequence space, adding to conventional analyses of driving forces which usually focus on binding energies. Indeed, an important finding of our work is that we have demonstrated how a genetic-algorithm framework that can alter the LLPS behaviour of proteins also enables us to probe the gradient in amino-acid sequence space directly. This can help us both to extend interpretations of why proteins that drive the formation of condensates might have evolved as they have and to gain greater control over intracellular LLPS.

We have coupled our genetic-algorithm technique to the coarsegrained model of Dignon *et al.*^33^ which is one of the best models currently available for probing the phase behaviour of protein solutions. This model has been validated against the singlemolecule experimental radii of gyration of a wide range of IDRs,^33^ is residue-specific, has been shown to reproduce well the experimental phase behaviour of various proteins under different conditions,^21,37,75–77^ and is computationally sufficiently inexpensive that it affords the determination of bulk LLPS properties for many sequences. Furthermore, the model accounts for key physicochemical aspects that determine the phase behaviour of proteins, such as the charge, size, relative hydrophobicity and flexibility of amino acids. However, as is the case with any coarse-grained model, it is still approximate and averages out other effects, in this case especially the specific contribution of π–π interactions, polarisation effects that give rise to cation–π interactions and the explicit role of water and ions in solution. The genetic algorithm we have introduced can of course straightforwardly be used with protein models of higher resolution and accuracy as they are developed, provided that sufficient computational resources are available.

Although the fine details of the phase behaviour we have observed may be model-specific, we have nevertheless shown two distinct driving regimes for enhancing or inhibiting LLPS to exist, namely hydrophobicity – including both the strength and range of relevant interactions – and charge patterning. In sequences such as the PLD of FUS, only one may be in operation, whilst in others, such as hnRNPA1-IDR, they may co-evolve, implying that both driving forces can contribute to LLPS simultaneously. Furthermore, in the hydrophobic driving regime in the case of the PLD of FUS, we have shown that there is a patterning length scale of 2-3 amino acids, which one can interpret in the context of the stickers-and-spacers model of proteins. Intriguingly, we have shown that although LAF1-IDR is charge-rich, charge patterning does not appear to co-evolve with hydrophobicity. In all cases studied, LLPS is facilitated by an increase in the mean size of the amino-acid residues of the proteins, which results in a more structured protein-rich phase, which in turn can favour condensate formation. It would be especially interesting to investigate in future work whether such a driving force for phase separation is more universal than might previously have been thought. Finally, we have demonstrated that the genetic-algorithm approach is successful at evolving sequences in the presence of other species in the medium; here, we have focussed on a mimic of RNA as a proof of concept. Significant changes to the gradient in sequence space are observed when another species is introduced into the system, indicating that our method is also suitable for investigating the co-evolution of proteins and for studying biologically relevant mixtures of different species. Since the effect of sequence modifications on the phase behaviour of many-component mixtures is much less intuitive to predict manually than it is in one-component systems, the ability to guide phase behaviour algorithmically is especially attractive.

In summary, we have presented a robust and powerful framework for systematically modulating the LLPS of proteins by evolving their amino-acid sequences. We have shown that the approach is able to provide direct insight into the nature of LLPS in protein solutions, demonstrating both what fundamental driving forces are in operation as well as providing specific guidance into the kinds of mutation that may help encourage or inhibit LLPS in practical applications. We have already drawn useful conclusions from the application of our approach to specific cases, contributing a significant piece of the puzzle towards a fuller understanding of the physical driving forces behind LLPS. As ever more accurate force fields of proteins in solution are developed, our approach promises to be particularly fruitful in furthering our understanding of the regulation of LLPS in biology, as well as representing a first step towards future engineering of phase-separating sequences.

## VI. METHODS

In the Supplementary material, we describe the coarse-grained potential, provide further details about the computational methods used, provide further analysis and additional supporting results, and provide the sequences of the proteins studied. Supporting raw data are available at the University of Cambridge Data Repository, doi:10.17863/cam.58351.

### Simulation methods

We performed molecular dynamics simulations of a coarsegrained implicit-solvent model of proteins^33^ in which each amino acid is represented as a bead. Neighbouring amino acids in a protein chain are connected by harmonic springs, while other beads interact with one another with a hydrophobicity-scaled Lennard-Jones (LJ) potential and a Debye-Hückel electrostatic potential. The model is discussed in more detail in Sec. SI-S1-a, and the simulation methods in Sec. SI-S1-b.

### Genetic algorithms

Genetic algorithms optimise properties of a system in ways inspired by biological adaptation of populations.^78–80^ All numerical parameters listed below were chosen to balance the need for high evolution speed due to an expensive fitness function and that sufficient diversity in the population be maintained to avoid premature convergence.

1. We define a chromosome of length *n*, **x**_i_ = (*x*_*i*1_, *X*_*i*2_,…*x_in_*), where in our case *x_ij_* is an amino acid. A set of *N* chromosomes defines an initial population *U*_0_ = {***x***_1_, ***x***_2_,…, ***x**_N_*}, where *U_t_* denotes the population of a given round *t*. The starting population in our case corresponds to mutated versions of the WT _***x***WT_ with mutation rate *P*, where *P* is the frequency at which an amino acid is exchanged for a random one picked from the natural set of 20. We use *P* = 0.01 in this work. A scalar fitness function *f (**x**)* denotes the property being optimised. We use the width of the phase diagram at a fixed temperature as a proxy for the critical solution temperature, and define the fitness function as

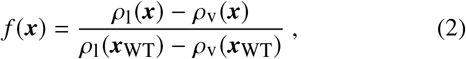

where *ρ*_1_ (***x***) is the average density of the ‘liquid-like’ protein-rich phase of sequence ***x*** and *ρ_v_ (**x**)* is the analogue for the ‘vapour-like’ protein-poor phase. We refer to individuals with high fitness value as ‘strong’ and to those with low fitness value as ‘weak’. In our case, *N* = 20. For genetic-algorithm runs in which the target is to reduce the critical solution temperature, we use as the fitness function the reciprocal of *f (**x**)*.
2. In each round *t*, we choose *N*_par_ = 8 parents *P* from the population, *P* ⊂*U_t_*. To achieve this, we use tournament selection:^80^ We first define *N*_par_ tournaments *T_i_*. Each tournament is a randomly drawn subset of *N*_tour_ elements from *U_t_*. The fittest sequence from each tournament becomes one of the parents. The tournament size *N*tour is therefore a direct scaling parameter governing the selection pressure. For our purposes, we have found that *N*_tour_ = 5 works well.
3. The parents are randomly divided into pairs (***a**, **b***) (where ***a*** ∈ ***P*** and ***b*** ∈ ***P***), and crossed over. Here, we swap sequences after a randomly chosen position *k* ∈ [1, *n*] in the sequences, such that

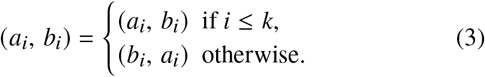
4. Another round of random mutations with frequency *p* is performed subsequently to cover previously unrepresented areas of sequence space.
5. The result from steps 3 and 4 is a set of children *C*, whose fitness is then evaluated. Children replace some chromosomes in the population. As fitness functions are relatively expensive to compute for our system, we use weak-population replacement,^78^ a greedy algorithm that can achieve rapid population evolution. Sequentially, each child ***c**_i_* ∈ *C* is compared to the weakest individual in the population, ***X***_weak_ ∈ *U*_t_. If *f* (***c**_i_*) > *f* (***x***_weak_), then ***c**_i_* replaces ***x***_weak_. The weakest individual may also be a previously inserted child.

Parallelisation can speed up genetic-algorithm progression.^81^ We use a simple master–slave approach since asynchronous schemes are not well suited to small populations; because all simulations are run for the same amount of wall-clock time, the overhead of the simple genetic-algorithm parallelisation employed is small compared to the duration of individual simulations.

## ACKNOWLEDGMENTS

We thank Jeetain Mittal and Gregory L. Dignon for their invaluable help with implementing their sequence-dependent coarsegrained protein model in Lammps. We acknowledge DiRAC funding from the Science and Technology Facilities Council. This project has received funding from the European Research Council (ERC) under the European Union Horizon 2020 research and innovation programme (grant agreement 803326). R. C.-G. is an Advanced Research Fellow from the Winton Programme for the Physics of Sustainability. A. G. is funded by an EPSRC studentship (EP/N509620/1). This work was performed using resources provided by the Cambridge Tier-2 system operated by the University of Cambridge Research Computing Service funded by the EPSRC Tier-2 capital grant EP/P020259/1.

## SUPPLEMENTARY MATERIAL

### S1 COMPUTATIONAL DETAILS

#### S1-a Coarse-grained model of proteins

The coarse-grained potential of proteins introduced by Dignon and co-workers^33^ is based on an amino-acid level description of protein chains. Each amino acid is represented by a bead. For amino acids that are covalently bonded in the protein of interest, their beads are connected by a harmonic spring,

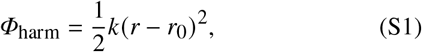

where *k* = 9.6 kcal mol^-1^Å^-2^ is the spring constant and *r*_0_ = 3.81 Å is the equilibrium bond length. Beads interact with one another through an Ashbaugh–Hatch modulated^82^ hydrophobicity-scaled (12,6)-Lennard-Jones (LJ) potential,

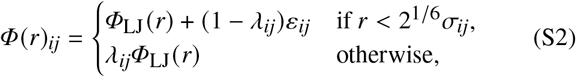

where *r* is the interparticle distance, and

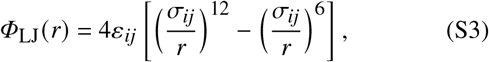

where *ε_ij_* is the minimum of the LJ potential, *σ_ij_* is the LJ diameter and *λ_ij_* is a hydrophobicity scaling parameter. In each case, *i* and *j* correspond to the amino acid types of the two particles. Residues that carry a charge (see Table S1) also interact with a Debye-screened^83^ electrostatic potential,

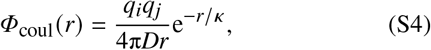

where *D* is the dielectric constant *D* = 80 *ε*_0_ (where *ε*_0_ is the electric constant), *κ* is the screening length (*κ* = 1 nm, corresponding to an ionic strength of 100 mm) and *q_i_* and *q_j_* are the charges of the amino acids.

In the work of Dignon and co-workers,^33^ two possibilities for assigning *λ_ij_*, *ε_ij_* and *σ_ij_* are presented. Here, we use the variant based on the Kim–Hummer model,^84^ which has been brought into the form of equation (S2) with parameters

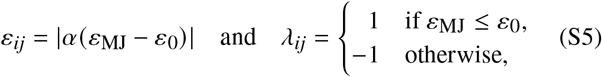

where *ε*_MJ_ is the Miyazawa–Jerningan empirical contact potential,^85^ and α = 0.228 and *ε*_0_ = 1 kcal mol^-1^ are benchmarked on experimental radii of gyration. The *σ_ij_* parameters are arithmetic means of effective Van der Waals radii of the amino acids, *σ_ij_* = (σ*_i_* + σ*_j_*)/2, which are given in Table S1.

A version of this model with a new parameter set^36^ was recently introduced, featuring a temperature dependence of the *λ_ij_* parameters to match experimental data more closely. We have not employed this potential here, as we are primarily interested in relative shifts of *T*_c_ and not its absolute value.

As an alternative to the Kim–Hummer model, a variation in the *λ_ij_* = (*λ_i_* + *λ_j_*)/2 parameters in an amino-acid specific way is also possible, while taking *ε_ij_* as a constant. This has been shown to yield comparable results.^33^ Whilst we do not employ this model here, we use these *λ_i_* values (Table S1) of the amino acids to quantify their hydrophobicity.

**TABLE S1.**
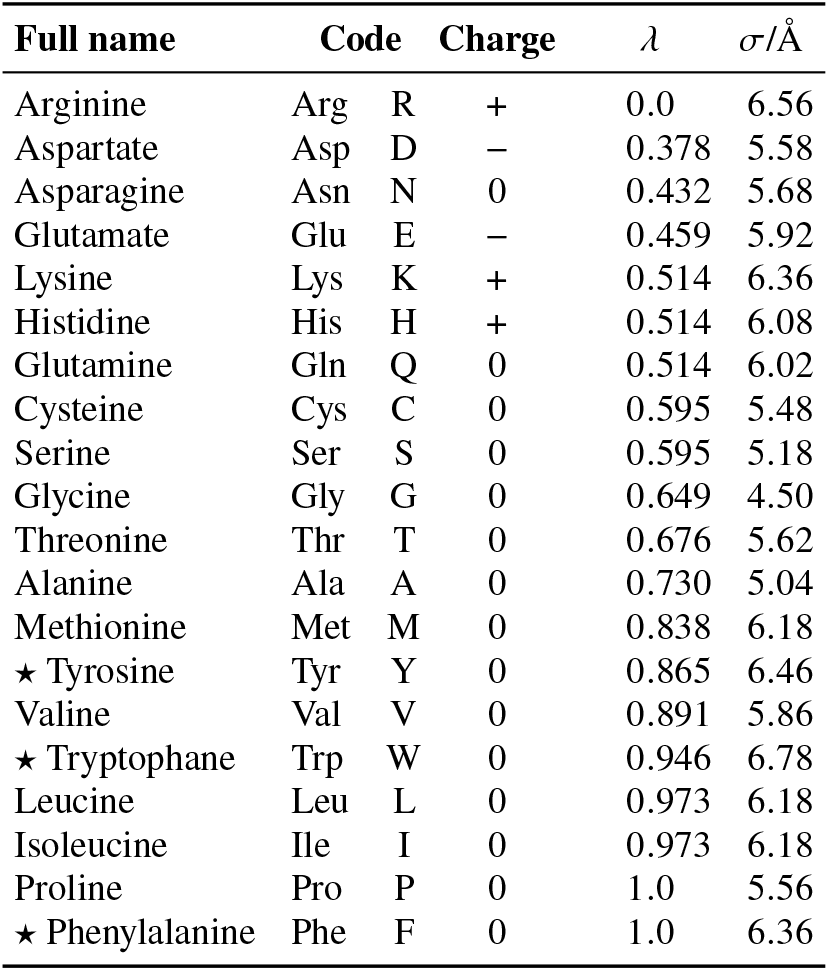
The 20 naturally occurring amino acids with their one- and three-letter codes, alongside their charges. Amino acids marked with a ‘✶’ are aromatic. The last columns give the *λ*- and σ-parameters which define the hydrophobicity scale.

#### S1-b Determining phase coexistence: simulation details

We studied LLPS with direct-coexistence molecular dynamics simulations^86^ in which a high- and a low-density phase coexist. Molecules can be exchanged between the two phases, allowing the densities to equilibrate across the interface [Fig. S1(a))]. In the limit of sufficiently large systems, where the interface becomes negligible compared to the bulk of each phase, this approach allows us to determine the densities of the two compositional phases. Direct-coexistence simulations provide an especially simple method of determining phase equilibria, particularly with liquid-like phases considered here. Since phase separation above the spinodal is a nucleation-initiated process, hysteresis may be a problem, and direct-coexistence simulations may require a careful calibration of the interface at the start of a simulation.^53^ However, with the kinds of coarse-grained potentials we are using, phase transitions are facile and an interface forms readily. Direct-coexistence simulations have therefore routinely been employed in computational studies of LLPS with such models.^87^

In order to construct a phase diagram, we perform direct-coexistence simulations at a number of temperatures [Fig. S1(a)]. We determine the densities of the coexisting phases by binning particles along the *z* axis and finding a least-squares best fit to two constant densities – with an interface of a system-specific width – to identify the low- and the high-density phases as well as the interfacial region [Fig. S1(b))]. Finally, the densities we extract are plotted on a temperature–density phase diagram, as for example in Fig. S2. To interpolate the data points close to the critical temperature, we use an empirical fit^30^ which, although it does not capture the behaviour of the system very well at low temperatures, is sufficient to find the approximate critical temperature and density. In particular, we fit simultaneously to

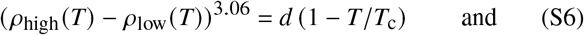

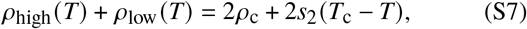

where ρ_high_ (*T*\*), ρ_low_ (*T*\*) and *ρ_c_* are the densities of the high-density and low-density phases and the critical density, respectively; *T*_c_ is the critical temperature and *d* and *s_2_* are fitting parameters. This works by numerically finding a best fit for the four constants - ρ_c_, *T*_c_, *d* and *s_2_* - from equations constrained by all pairs of Phigh (*T)* and ρ_low_ (*T*) deemed by inspection of the curvature of the observed data series to lie below *T*_c_.

We note that in direct-coexistence simulations, above the critical temperature, the system forms a single supercritical fluid; the density fitting outlined above, which assumes two distinct phases have formed, will thus produce non-physical results reflecting the natural density fluctuations of the system; any densities so determined do not correspond to coexisting phases. Nevertheless, such spurious points help us to ascertain that we have already crossed the critical point, forming a characteristic ‘protrusion’ of the coexistence region towards higher temperatures. The presence of such features at the appropriate point can thus provide an additional check that we have determined the critical temperature correctly, and so we have included them in phase diagrams as greyed-out points for reference.

**FIG. S1.**
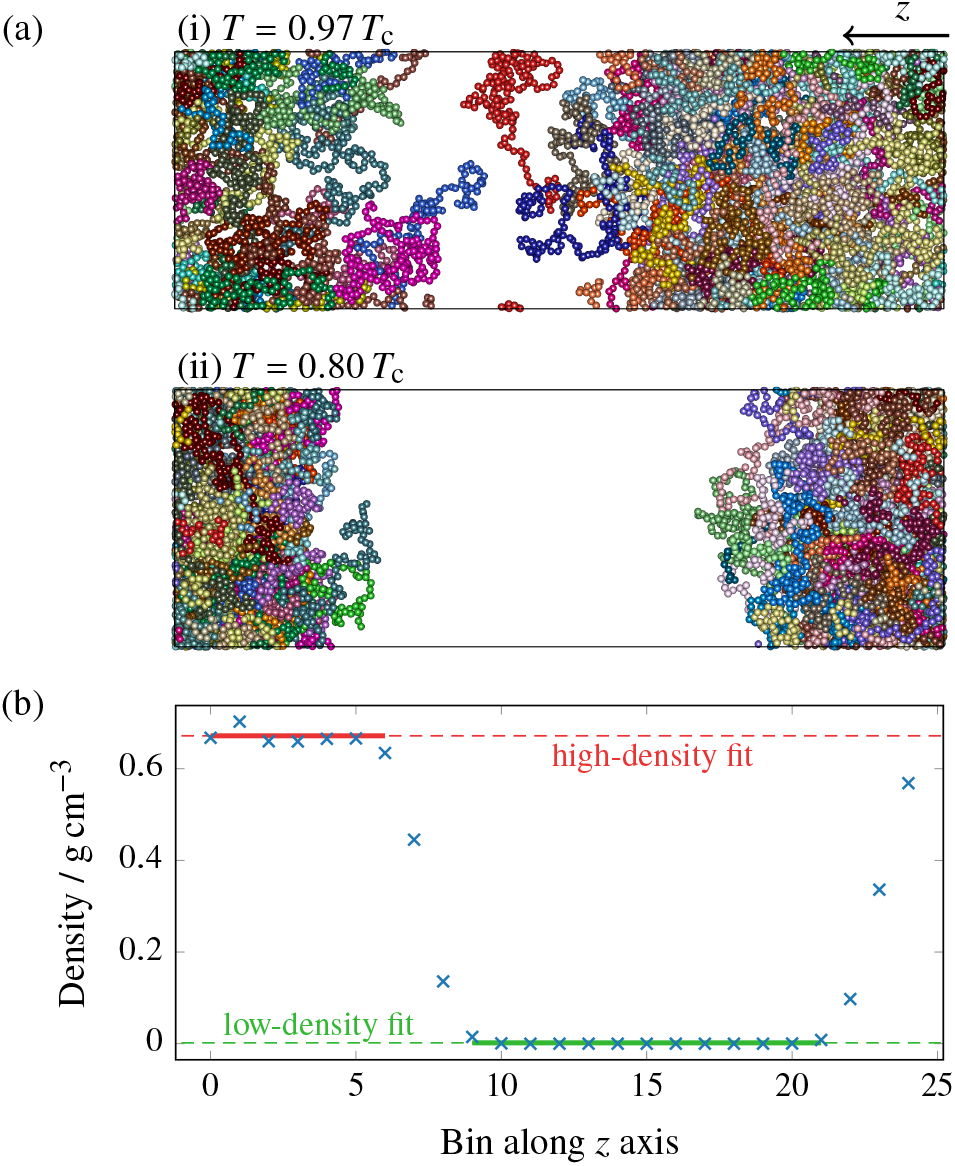
(a) Orthographic projections of simulation boxes of FUS at two temperatures below *T*_c_, as indicated, where different colours represent different protein chains. The box is periodic in all directions. (b) Example of a fitted density profile for FUS at *T* = 0.8 *T*_c_. The simulation box along the *z* direction is split into 25 bins and the average density is computed within each bin, indicated by blue crosses. Points from the regions where solid lines are shown were used to compute a fit to a constant for each of the high- and the low-density phases, taking into account an interface of finite thickness.

We performed molecular dynamics simulations with the Lammps simulation package,^88,89^ using a velocity Verlet integrator with a time step of δ*t* = 10 fs. Direct-coexistence simulations were run in the canonical ensemble with a Langevin thermostat^90^ with a damping time of 10^4^ *δt*. We used a tetragonal simulation box with periodic boundary conditions, with typical dimensions 134.4Å × 134.4Å × 403.2Å with 64 chains for FUS and hnRNPA1, and 267.2 Å × 267.2 Å × 1336.2 Å with 512 chains for LAF1.

### S2. HYDROPHOBIC SEQUENCE OPTIMUM

**FIG. S2.**
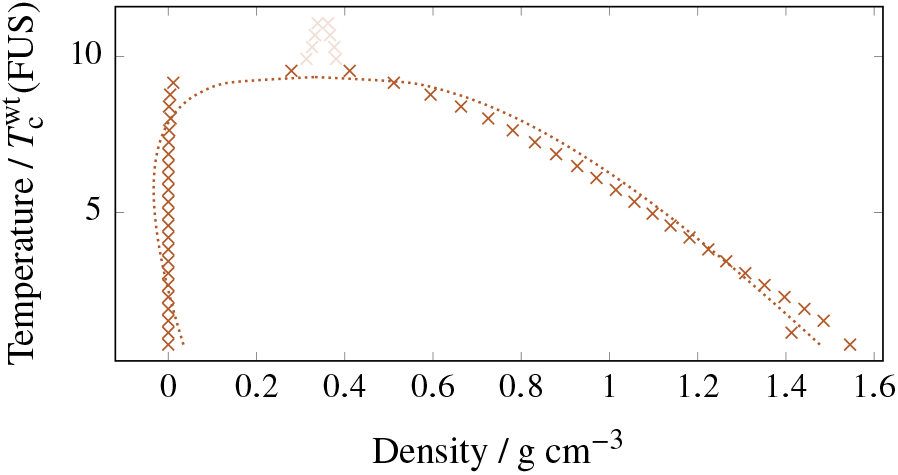
Phase diagram of the (Phe)163 sequence, the same length as FUS-WT. The critical temperature is approximately nine times that of FUS-WT. The dotted line is a fit, and greyed-out points lie above the critical point, as detailed in Section S1-b.

Figure S2 shows the phase diagram of a possible optimum of hydrophobicity-driven LLPS within the CG model. While assessing an optimum of the charge-driven case is difficult, it possible to make a guess about the scaled LJ potential by inspection of the model parameters. The highest *ε_ij_* value is for a Phe-Phe interaction, which is also longer-ranged than for most other amino acids. The phase diagram shown illustrates how large the changes in critical temperature can be, but it also demonstrates that trivially optimising the critical temperature results in a biologically uninteresting sequence.

### S3. MINIMISING THE CRITICAL TEMPERATURE

**FIG. S3.**
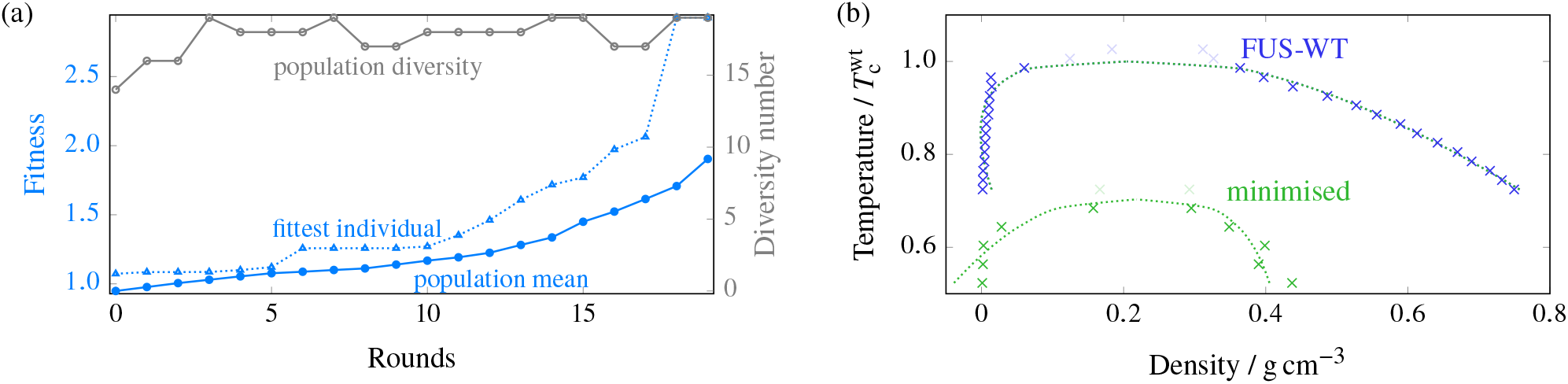
(a) Typical GA progression for FUS where the fitness function *minimises* the upper critical solution temperature. The fitness function [defined as the reciprocal of Eq. (2) of the main manuscript] increases by ~90 % over 20 rounds. The fittest individual can be considerably fitter than the mean. The population diversity, i.e. the number of distinct sequences present in the overall population of 20, is generally very high. (b) Comparison of representative phase diagrams before and after genetic-algorithm runs, confirming that the fitness function choice was suitable. Dotted lines are fits, and greyed-out points lie above the critical point, as detailed in Section S1-b.

### S4. LOCAL STRUCTURE OF THE LIQUID-LIKE PHASE

**FIG. S4.**
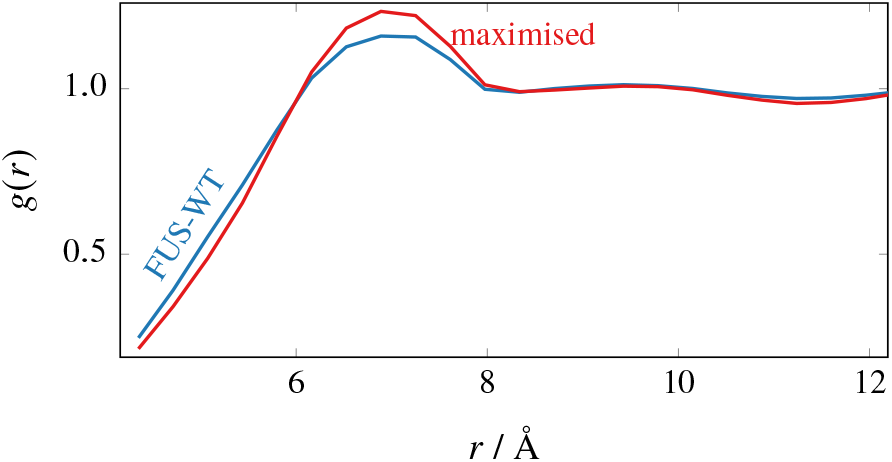
Comparison of the radial distribution function *g(r)* before and after maximising the phase diagram width of FUS. The data were computed at the same number density *N/V* = 6.757 nm^-3^ and temperature 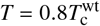. The temperature chosen is below the critical point in both cases, and the density in both cases corresponds to the protein-rich (liquid-like) phase, i.e. a point that lies above the binodal line on the phase diagram. The radial distribution function is computed by finding for each bead *i* in the system the number of all non-harmonically bonded other beads within a distance *r* + *δ_r_* of the each other for bins of width δ*r*, averaging over each bead *i* in the system, and normalising the result by the volume element and the (common) number density. In the case of the evolved system, the more pronounced nearest-neighbour maximum indicates the local environment is more structured than in the case of WT-FUS.

### S5. CHUNK SHUFFLING EXAMPLE RUNS

**FIG. S5.**
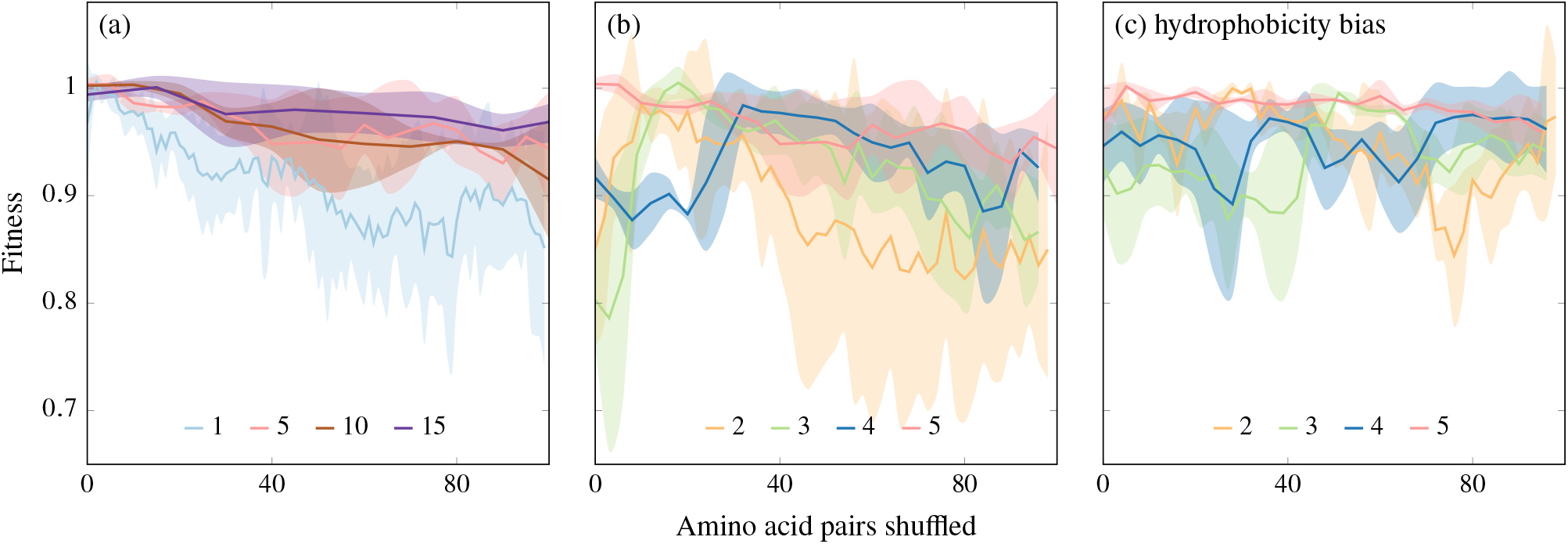
(a) Chunk shuffling at chunk lengths 1,5, 10 and 15. Chunk length 1 lies significantly below the other curves. Shaded areas are standard deviations from 3 (6 for chunk length 1 and 5) shuffling runs. (b) Chunk shuffling with focus on chunk lengths around a length scale of 2-3 amino acids. Shaded areas are standard deviations from 3 (6 for chunk lengths 2 and 5) shuffling runs. (c) Hydrophobicity-biassed chunk shuffling, only allowing exchanges between the top 30 % chunks by hydrophobicity. Shaded areas are standard deviations from 3 shuffling runs.

### S6. OPTIMISATION AND PHASE DIAGRAM OF HNRNPA1

**FIG. S6.**
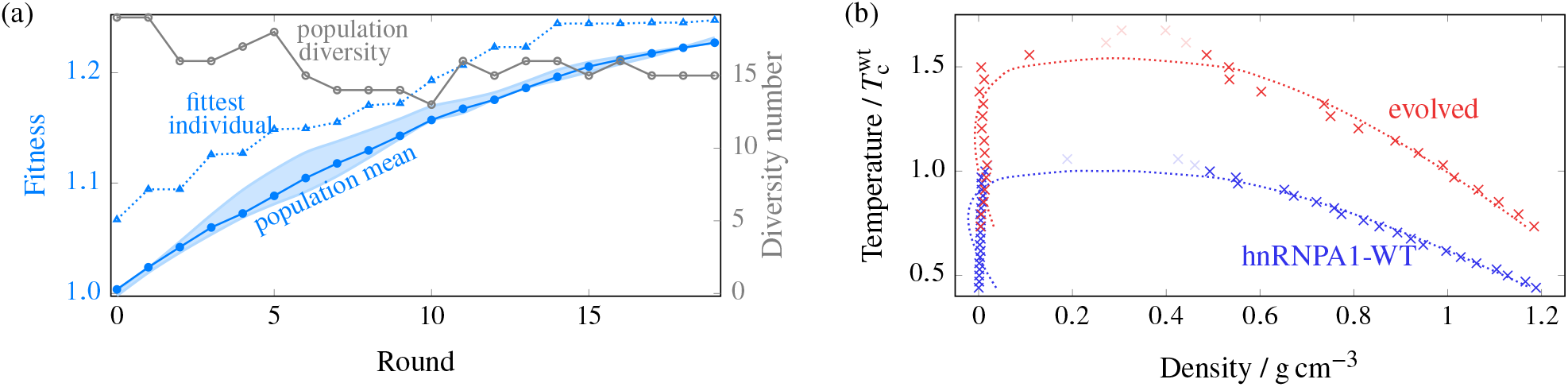
(a) Typical GA progression for hnRNPA1. The fitness function [Eq. (2) of the main manuscript] was evaluated at 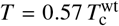, and increases by ~20 % over 20 rounds, while maintaining population diversity. The shaded area corresponds to the range of values of the mean fitness obtained from 3 independent genetic-algorithm runs. (b) Comparison of representative phase diagrams before and after genetic-algorithm runs, showing a ~50 % increase in critical temperature. Dotted lines are fits, and greyed-out points lie above the critical point, as detailed in Section S1-b.

### S7. OPTIMISATION AND PHASE DIAGRAM OF LAF1

**FIG. S7.**
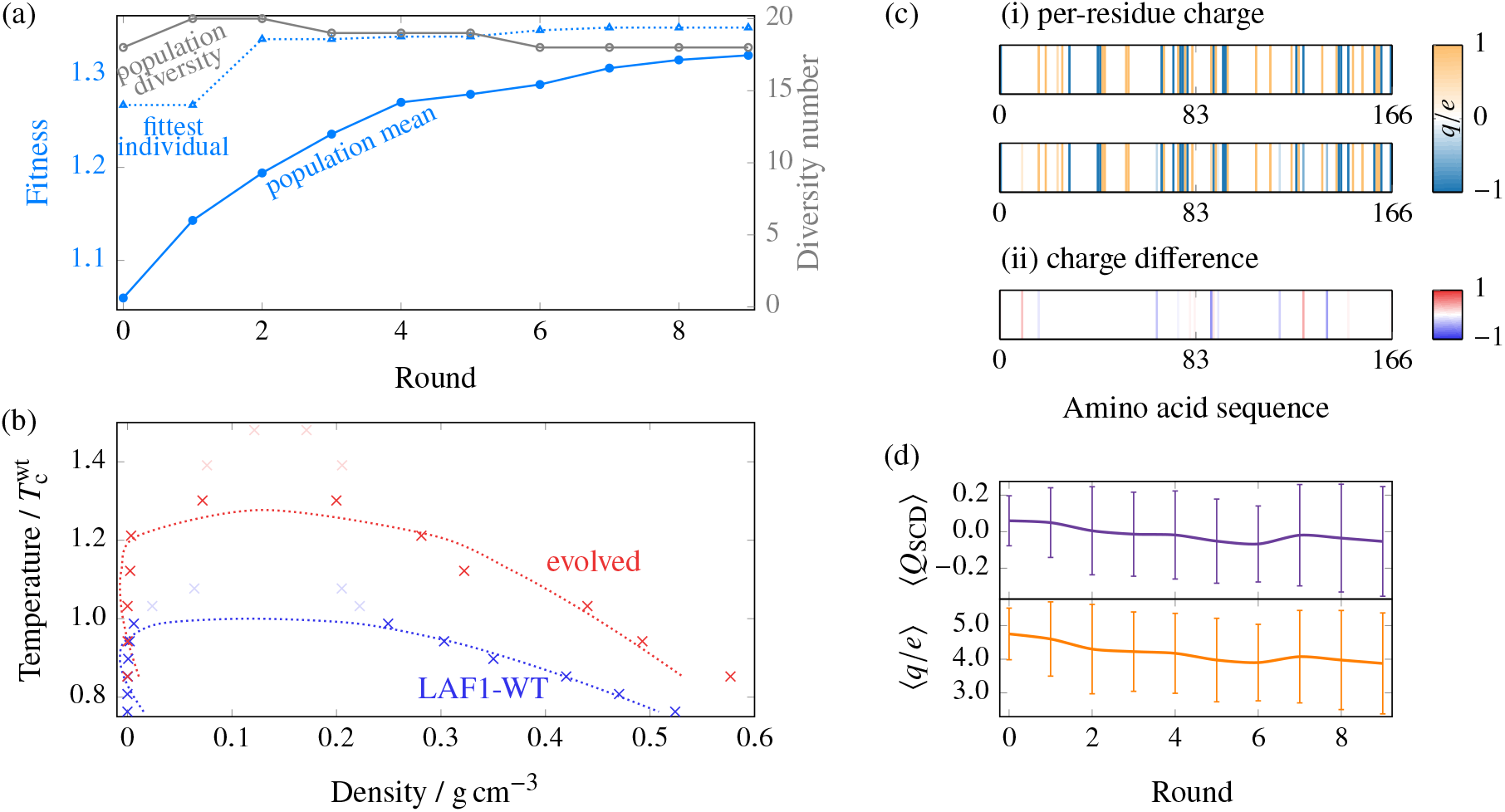
(a) Typical GA progression for LAF1. The fitness function [Eq. (2) of the main manuscript] was evaluated at 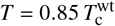, and increases by ~30 % over 9 rounds, while maintaining population diversity. To obtain meaningful density data in the light of thick interfaces, the simulation box had to be doubled in all directions compared to simulations of FUS and hnRNPA1. (b) Comparison of representative phase diagrams before and after genetic-algorithm runs. Although the phase diagram close to the critical point is especially difficult to equilibrate because of interfacial effects in this system, data points at lower temperatures suggest that the critical temperature increases by ~30 % by the end of the GA optimisation. (c) Map of (i) per-residue charges along the sequence for WT and evolved LAF1 and (ii) the difference at the end of the genetic-algorithm run. Partial charges reflect only partial carriage in the population. More charges are lost than created during the course of the genetic-algorithm runs. (d) The sequence charge decoration (SCD) and the charge number (*q/e*) of the population over the course of the genetic-algorithm run.

### S8 LIQUID CHARACTER OF PHASES

**FIG. S8.**
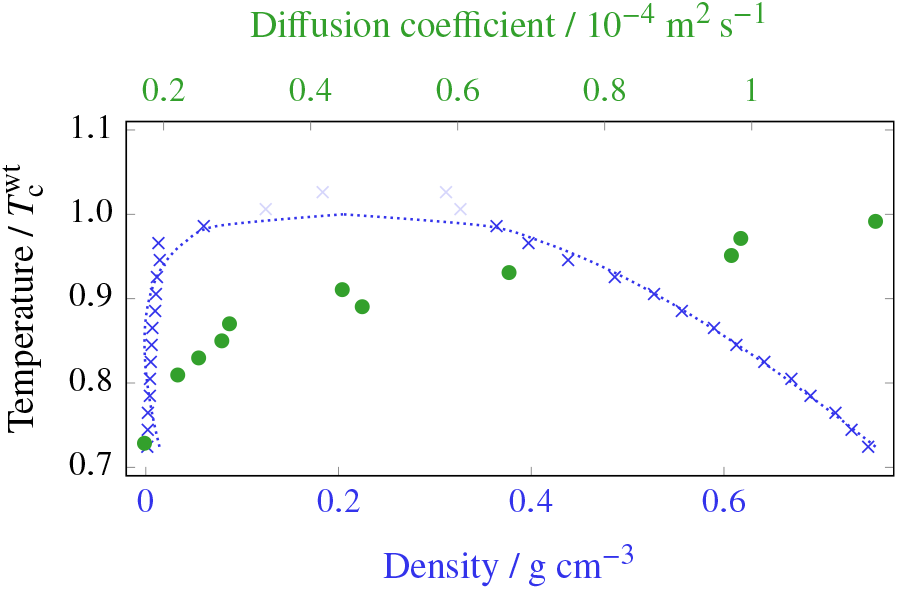
Phase diagram of wild-type FUS (in blue) alongside diffusion coefficients for the liquid-like phase (in green). The densities of the coexisting phases were first determined in direct-coexistence simulations at a range of temperatures. The diffusion coefficient was then computed in a canonical-ensemble simulation at the density corresponding to the liquid-like (high-density) phase. A non-linear relationship exists between the diffusion coefficient and temperature, but there is no indication of a discontinuous change that might indicate a glass transition.

When studying LLPS, it is important to control the liquid character of the observed phases, as proteins can also undergo gelation^49^ and glass transitions.^91^ While the study of general polymer dynamics in the CG model is not the objective of this work, and dynamical properties of coarse-grained models do not usually faithfully reproduce real dynamics, we nevertheless computed the per-bead diffusion coefficient for our systems to ascertain that a glass transition has not occurred. Ballistic and sub-diffusive regimes may precede the diffusive regime, and in polymer systems, several sub-diffusive regimes can often be observed.^92,93^ The diffusion coefficient *D* can be obtained by the Einstein relation^94^ for the mean squared displacement, 〈Δ*r*^2^〉 = *6D_t_*, which holds in the diffusive regime of *long* times *t*. We have computed diffusion coefficients in this way over a range of temperatures, and show these alongside the phase diagram of FUS-WT in Fig. S8. The diffusion coefficients shown are only qualitative, in the sense that in the potential we use, many degrees of freedom have been coarse-grained away, and the unit of time is not directly comparable to experiment. However, ratios of diffusion coefficients are nevertheless meaningful. The temperature dependence of the diffusion coefficients is non-linear and the presence of significant noise makes a detailed analysis difficult; however, there is no obvious discontinuous change in the diffusion coefficient as a function of temperature in these data even at low temperatures in the liquid phase, which we take as justification that our simulations describe LLPS rather than glass formation, which could complicate our interpretation of the results.

### S9 AMINO-ACID SEQUENCES OF PROTEINS STUDIED

We give below the amino-acid sequences of the prion-like IDR of FUS, hnRNPA1-IDR and LAF1-IDR^29^ studied in this work, using one-letter codes [Table S1] for the amino acids.

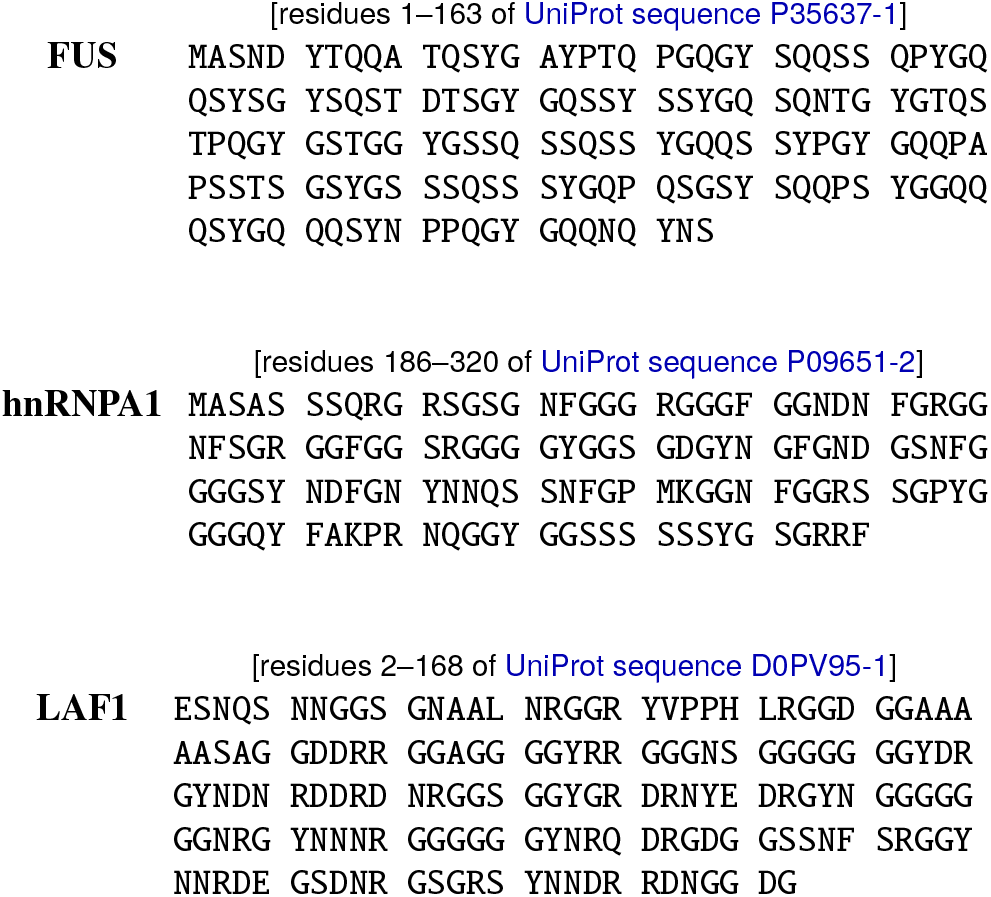

### S10 EVOLVED SEQUENCES

Below, we provide example output sequences, taken from the final populations of the appropriate GA runs as described in this work, using one-letter codes [Table S1]. The residues that have changed compared to the initial sequence are highlighted in red. Sequences of full populations as a function of the progression of all our GA runs can be found in the supporting data.

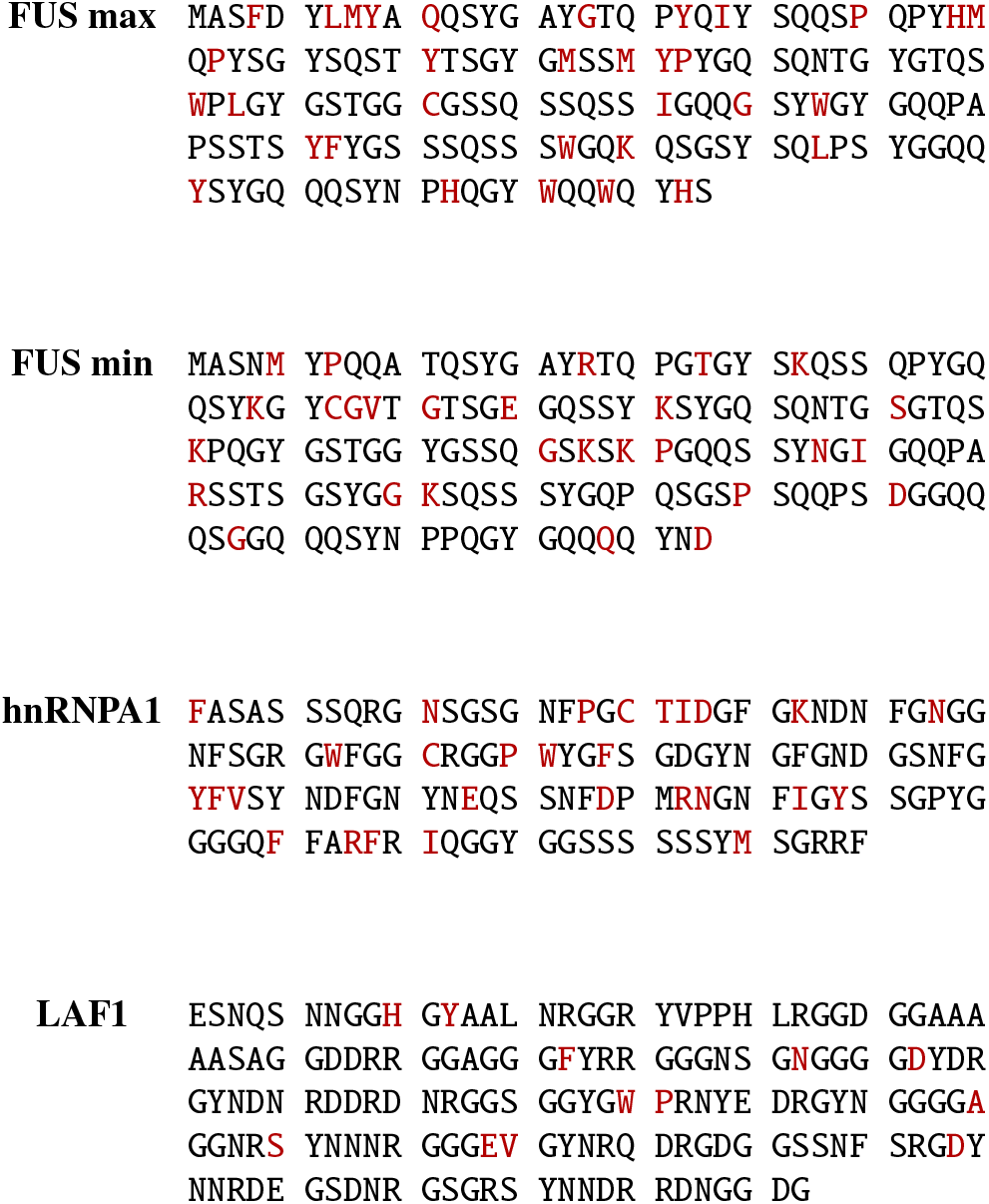

## Notes

### Competing Interest Statement

The authors have declared no competing interest.

